# Fourier-transform-based attribution priors improve the interpretability and stability of deep learning models for genomics

**DOI:** 10.1101/2020.06.11.147272

**Authors:** Alex M. Tseng, Avanti Shrikumar, Anshul Kundaje

## Abstract

Deep learning models can accurately map genomic DNA sequences to associated functional molecular readouts such as protein–DNA binding data. Base-resolution importance (i.e. “attribution”) scores inferred from these models can highlight predictive sequence motifs and syntax. Unfortunately, these models are prone to overfitting and are sensitive to random initializations, often resulting in noisy and irreproducible attributions that obfuscate underlying motifs. To address these shortcomings, we propose a novel attribution prior, where the Fourier transform of input-level attribution scores are computed at training-time, and high-frequency components of the Fourier spectrum are penalized. We evaluate different model architectures with and without attribution priors trained on genome-wide binary or continuous molecular profiles. We show that our attribution prior dramatically improves models’ stability, interpretability, and performance on held-out data, especially when training data is severely limited. Our attribution prior also allows models to identify biologically meaningful sequence motifs more sensitively and precisely within individual regulatory elements. The prior is agnostic to the model architecture or predicted experimental assay, yet provides similar gains across all experiments. This work represents an important advancement in improving the reliability of deep learning models for deciphering the regulatory code of the genome.

## 1 Introduction

Transcription factors (TFs) are proteins that regulate gene activity with extraordinary specificity by recognizing and binding to short DNA sequence patterns—or “motifs”—in regulatory elements in the genome. High-throughput experiments have been used to profile genome-wide regulatory activity in diverse cell types and tissues [1]. Chromatin immunoprecipitation sequencing experiments (e.g. ChIP-seq, ChIP-exo, and ChIP-nexus) provide binding readouts of specific targeted TFs at each base (DNA letter) in the genome [1]. Chromatin accessibility experiments (e.g. DNase-seq and ATAC-seq) provide aggregate readouts of all protein–DNA contacts at each base in the genome [1]. Deep neural networks—particularly convolutional neural networks (CNNs)—have achieved state-of-the-art performance in mapping DNA sequence to TF binding and chromatin accessibility profiles [2, 3, 4]. These models accept fixed length DNA sequence segments of the genome as an input, and—through a series of convolutional layers—predict molecular labels as measured by a regulatory profiling experiment. A common goal of these genomic deep learning models is to ultimately identify the regulatory sequence code (i.e. motifs and their syntax) underpinning genomic regulation, and this has motivated the development of a wide suite of tools to infer base-pair-resolution importance (i.e. “attribution”) scores in the input DNA sequences to reveal the regulatory code [5, 6, 7, 4].

Unfortunately, the high capacity of these models make them more prone to overfitting on training set noise rather than true signal [8]. Querying such models for their attribution scores often reveals their reliance on regions of the input sequence that are not biologically relevant to the task at hand. Perhaps worse, their attributions are irreproducible and highly sensitive to random initializations (Figure 1). This poses a hindrance to the reliable identification of motifs driving genomic regulation from the attribution scores.

**Figure 1:**
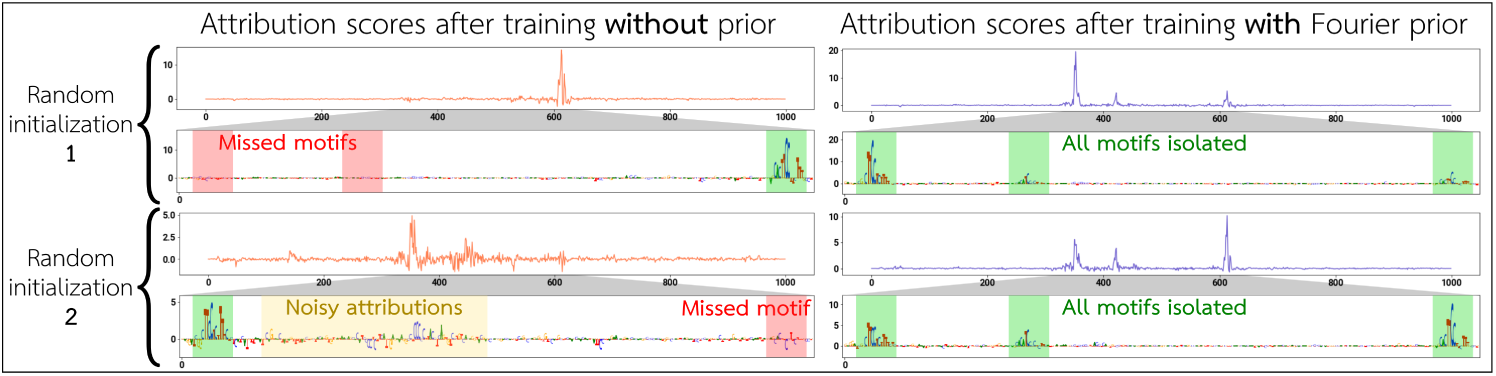
Models trained with the standard approach (left) irreproducibly miss motifs in the underlying sequence and noisily rely on irrelevant regions of the input. When training with the Fourier-based attribution prior (right), models consistently and cleanly identify the driving motifs. The examples shown are from binary binding models of the SPI1 TF from TF ChIP-seq experiments [1]. See Supplementary Figures S14–S15 for more examples.

There is a need to improve the interpretation of high-capacity genomic deep learning models with the goal of downstream motif discovery. To address this need, we propose a novel attribution prior [9, 10] based on Fourier transforms to directly reward the notion of interpretability during model training. That is, at training time, we impose a secondary loss function that penalizes the network for improper attributions, thereby *explicitly* training the model to maximize interpretability along with correctness of predictions. We show that our Fourier-based attribution prior can be flexibly applied to different model architectures using a diverse set of real experimental data, and offers improvements in interpretability, motif discovery, and learning stability (Figure 1).

## 2 An attribution prior for genomics based on Fourier transforms

### 2.1 Formulation of the Fourier-based attribution prior

Regulatory DNA sequences are sparsely composed of short functional sequence motifs (*∼* 6–20 bases in length) with soft syntactic constraints on motif combinations, density, spacing and orientation. To maximize interpretability with the goal of motif recovery, CNNs trained on regulatory data would ideally place importance (i.e. attribution) only on informative motifs, and little importance on the irrelevant background sequence. Let *x* represent a one-hot encoded input sequence to the model, *y* represent the true labels, and *f* represent the model prediction function. Let *g*(*x, f*) represent the per-position attribution scores on the input *x*, where *g*(*x, f*) is a vector of attributions of length equal to the length of *x*. We train our models with a two-part loss function: *L*(*x, y, f*) = *L*_*c*_(*f* (*x*), *y*) + *λL*_*p*_(*g*(*x, f*)), where *L*_*c*_ is the standard correctness loss for the model, and *L*_*p*_ is the attribution prior loss. The choice of *g* depends on the model type (Supplementary Methods Sec. 2.3). In this work, we formulate *g* using input gradients for computational efficiency, and subsequently show that the improvements granted by the prior follow through to an alternative reference-based attribution method.

Our Fourier-based prior loss is computed by taking the Fourier transform of the attribution vector *g*(*x, f*). Let *m* denote the magnitudes of the positive-frequency Fourier components of (a slightly smoothed version of) *g*(*x, f*), and 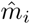 denote the *i*^th^ component of 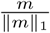 (smaller *i* correspond to lower frequencies) (Supplementary Figure S1).

We penalize the high-frequency components in 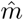 as follows:

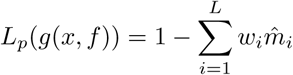

Where

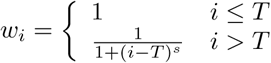

This attribution prior penalizes the model for attempting to place importance along the input DNA sequence in bursts shorter than the limit set by *T*, which denotes the *minimum* length of sequence that can be considered a reasonable motif. This limit is softened by the parameter *s*, which controls the rate at which this penalty is smoothly reduced for higher frequencies. *T* is set based on prior knowledge by assuming no motif will be shorter than 7 bp, while *s* is fixed to 0.2 for all models (Supplementary Methods Sec. 2.3). Note that due to the *ℓ*_1_-normalization of 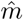, the value of *L*_*p*_ is bounded between 0 and 1. Fourier transforms are linear and surprisingly fast to calculate, resulting in an attribution prior *L*_*p*_ that is efficient to compute and differentiate. Furthermore, by using Fourier transforms, our prior is effectively invariant to the locations and the number of motifs within an input sequence.

### 2.2 Training data and model architectures

To demonstrate the flexibility of the Fourier-based prior, we train on four different experimental datasets over two different model architectures. Our experimental datasets target the SPI1 protein (TF ChIP-seq) in 4 cell types [1]; the GATA2 protein (TF ChIP-seq) in 3 cell types [1]; the Nanog, Oct4, and Sox2 proteins (TF ChIP-seq) in mouse embryonic stem cells [4]; and chromatin accessibility (DNase-seq) in the K562 leukemia cell line [1]. These datasets vary not only in the type of experimental assay being predicted, but also in the complexity of motif syntax driving the measurements. For each dataset, we train both binary models and profile models. Binary models predict a binary label for each 1000-base-pair-long input sequence (i.e. whether or not there is a statistically significant peak in the TF ChIP-seq or DNase-seq accessibility signal overlapping the mid point of the sequence [1]) (Supplementary Figure S2). Profile models, however, predict base-resolution TF ChIP-seq or DNase-seq profiles from each 1346-base-pair (bp) input sequence, and therefore are able to finely track motifs based on associated patterns in the shape of the output profile (Supplementary Figure S3) [4].

When we train binary models with the Fourier-based prior, the prior loss weight *λ* is set to 1. On profile models, the prior loss weight is selected to be half of the value of the correctness loss on a model trained without the prior. In both architectures, this ensures that the value of the prior loss is on a similar scale as the correctness loss (Supplementary Methods Sec. 2.3, Supplementary Figure S4).

## 3 Fourier-based priors improve signal-to-noise ratio for the detection of predictive motifs

We consider models trained with and without the Fourier-based prior, and compare the interpretability of the models using DeepSHAP [11] scores, which extend DeepLIFT [7] backpropagation-based attributions using Shapely values. While the attribution prior computed at training time operated on input gradients due to efficiency, we rely on DeepSHAP attribution scores to assess a model’s interpretability because DeepSHAP scores provide base-pair-level attributions against a biologically meaningful background—in this case, shuffled versions of the input sequence that preserve dinucleotide frequencies (as recommended by Shrikumar et al. [7]).

Over all of our datasets and model architectures, we find that training with the Fourier-based prior significantly denoises attribution scores and dramatically improves the detection of predictive motifs near ChIP-seq/DNase-seq peak summits (local maxima in the profiles around which driver motifs are expected to be found). A model that elevates attributions only at the motifs—and has flat attributions outside motif regions—is expected to have a reduction in the high-frequency Fourier components and the Shannon entropy of the input sequence attributions. We see a reduction in both metrics on all of our datasets and architectures (Supplementary Table S1, Supplementary Figure S5). Using a Wilcoxon rank test, all improvements in test-set interpretability when training with the Fourier-based prior are significant at the 1 × 10^−6^ level.

We visually show the improvement in interpretability over examples of peak sequences in the test set, including the ability of models trained with the Fourier-based prior to cleanly highlight motifs (Figure 2, Supplementary Figure S6).

**Figure 2:**
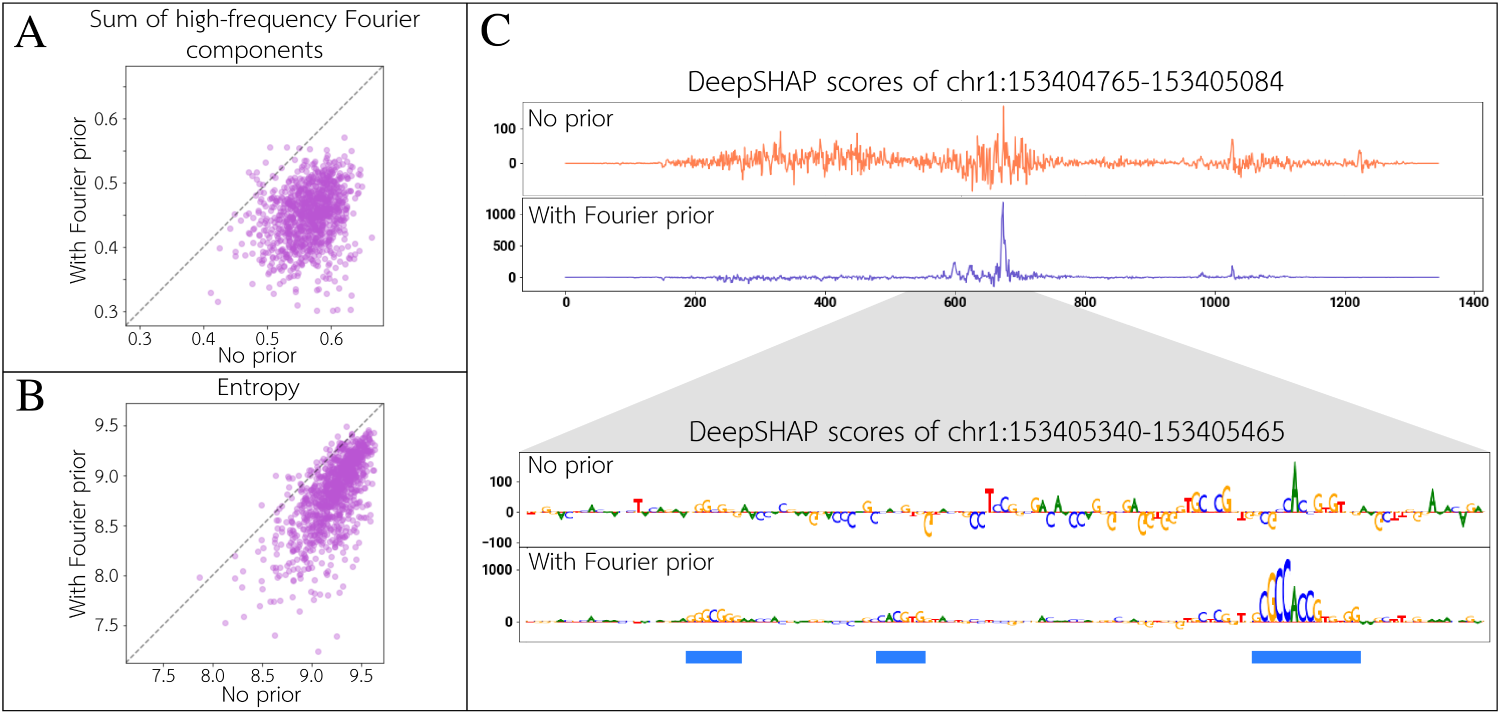
On K562 DNase-seq profile models, for each sampled test-set sequence, we compare the: **A)** sum of high-frequency normalized Fourier components; and **B)** Shannon entropy of the DeepSHAP attributions between models trained with versus without the Fourier-based prior. **C)** At a particular K562 open chromatin peak, we show the attributions across the entire input sequence, and the base-pair-level attributions around the summit region. The model trained with the Fourier-based prior cleanly highlights 3 motifs centered around the peak summit, matching relevant transcription factors (left to right: SP1, CLOCK, and CTCF).

## 4 Fourier-based priors improve sensitivity and specificity of motifs

On multi-task models trained to predict binding of the 3 TFs, Nanog, Oct4, and Sox2, we perform motif discovery and motif instance calling, and compare the calls to independently collected gold-standard ChIP-nexus experiments [4] that can highlight putative bound motifs at very high resolution (*∼* 10 bp). We use TF-MoDISco, a motif discovery approach, to distill recurring TF-binding motifs across multiple input sequences from base-resolution importance scores [6]. TF-MoDISco first finds predictive subsequences of high importance (called “seqlets”) across all input sequences. Seqlets are subsequently aligned, clustered, and summarized into a non-redundant set of motif patterns. We then use these motifs to scan test-set peak sequences and their attributions to call high confidence matches to the motif (i.e. motif instances). For both model architectures, we find that attributions derived from the models trained with the Fourier-based prior improve the quality of motifs discovered by TF-MoDISco, as well as the sensitivity and specificity of called motif instances supported by ChIP-nexus peaks (Supplementary Figures S7–S9).

We show an example of the improvement in motif discovery and motif calls using the multi-task Nanog/Oct4/Sox2 profile models. Specifically, on the Nanog prediction task, TF-MoDISco identified the known ATCAA and GGAAAT Nanog motifs (among others), in addition to the Oct4-Sox2 motif [4]; the Nanog motifs were missed without the prior (Figure 3). Additionally, motif instances called using discovered motifs with the prior show substantially improved support from independent high-resolution ChIP-nexus data.

**Figure 3:**
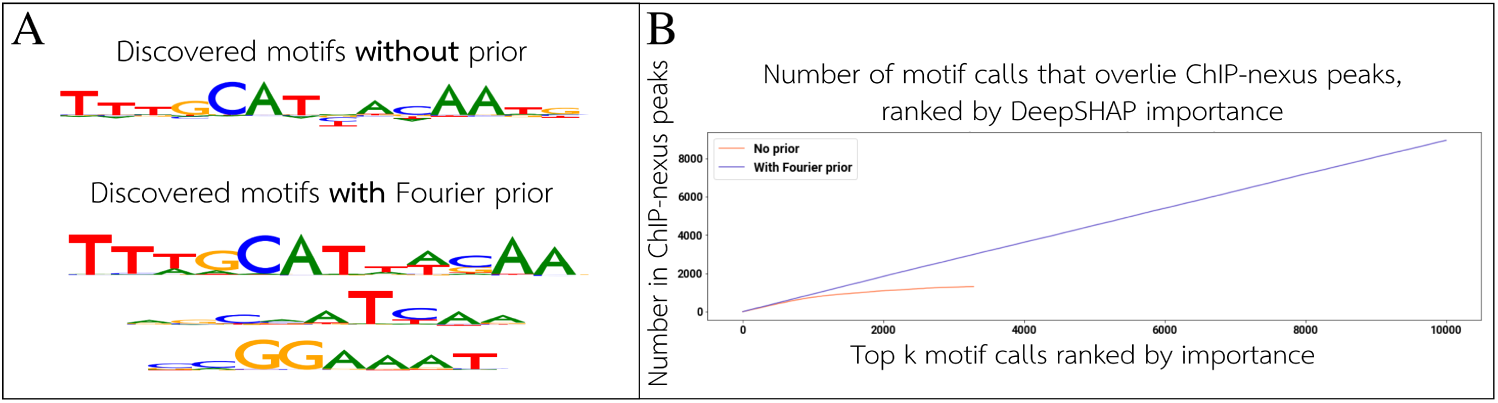
**A)** We show a subset of relevant motifs for the Nanog binding prediction task discovered by TF-MoDISco when training with versus without the Fourier-based prior. The ATCAA and GGAAAT Nanog motifs [4] are missed by the model trained without the prior. **B)** Using the motifs discovered by TF-MoDISco, we call motif instances on test-set peak sequences. Ranked by total attribution magnitude, we compute a cumulative count of motif calls that overlap gold-standard bound motif instances from independent high-resolution Nanog ChIP-nexus data. The prior substantially improves the recall of gold-standard motif instances.

## 5 Fourier-based priors improve prediction performance of binary models

Whereas profile models learn to predict continuous ChIP-seq or DNase-seq read coverage profiles at base-resolution, the more popular models trained on lower-resolution binary labels (e.g. bound vs. unbound) do not benefit from profile shape information, and thus are particularly prone to overfitting [4]. On all of our binary models, we see improvement in validation and test set performance when training with the Fourier-based prior (Supplementary Figure S11). Table 1 shows the best validation and test set performance achieved when training with versus without the Fourier-based prior, over 30 random initializations each.

**Table 1:** Binary model performance on test set

|  | Test accuracy |  | Test auROC |  | Test auPRC |  |
| --- | --- | --- | --- | --- | --- | --- |
|  | No prior | With prior | No prior | With prior | No prior | With prior |
| SPI1 | 87.996% | **88.347%** | 0.954 | **0.956** | 0.853 | **0.858** |
| GATA2 | 85.519% | **86.216%** | 0.924 | **0.928** | 0.733 | **0.745** |
| Nanog/Oct4/Sox2 | 84.640% | **85.437%** | 0.906 | **0.908** | 0.655 | **0.661** |
| K562 | 90.049% | **90.362%** | 0.966 | **0.968** | 0.963 | **0.966** |

We also show a more detailed depiction of the distribution of performance for binary models trained to predict SPI1 binding (Figure 4).

**Figure 4:**
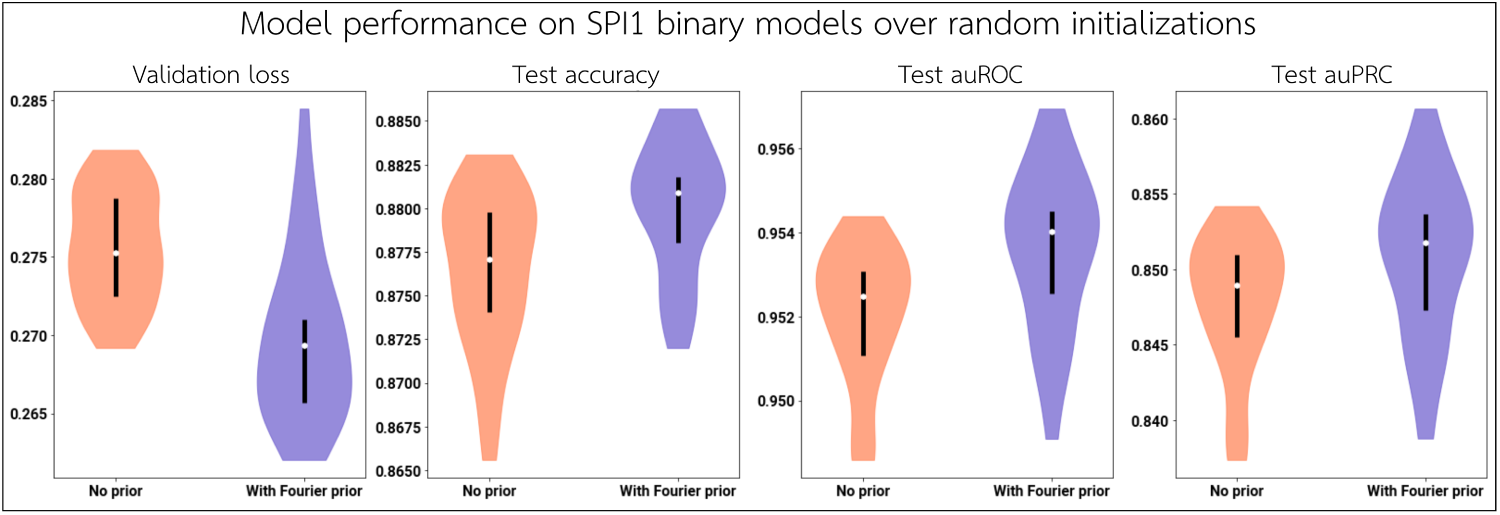
We train binary models to predict SPI1 binding over 30 random initializations without the prior, and another 30 with the Fourier-based prior. We show the distribution of the validation loss, test-set accuracy, test-set auROC, and test-set auPRC across the 30 random initializations in each condition.

Training on only 1% of the training data reveals a similar trend, showing that the Fourier-based prior assists in improving model generalizability even when the model is trained on sparse data (Supplementary Figure S12).

## 6 Fourier-based priors improve stability of attribution scores

Over all of our datasets and model architectures, we find a dramatic improvement in the stability of attributions when models are trained with the Fourier-based prior. For each dataset, on a set of test-set peak sequences, we compute the similarity of DeepSHAP attribution scores between the models to quantify how consistently the models learned on each particular sequence. We employ continuous Jaccard similarity—a metric used by TF-MoDISco—as it is designed for comparing similarity between two importance score tracks, accounting for similarity of both sequence and attribution scores [6] (Supplementary Methods Sec. 7).

The Fourier-based prior improves the stability of the attribution scores across random initializations, thereby allowing robust inference of motifs (Table 2; Supplementary Figures S13–S15). The improvements in learning stability are even stronger between models trained on significantly smaller training sets (i.e. 1% of the original set) versus the entire training set (Table 2; Supplementary Figures S13, S16). This suggests that the models trained on sparse data learn more similarly to when they are trained with the full dataset. Using a Wilcoxon rank test, all improvements in test-set stability when training with the Fourier-based prior are significant at the 1 × 10^−6^ level.

**Table 2:** Learning stability on test-set sequences (continuous Jaccard similarity)

|  |  | Similarity across random initializations |  | Similarity between training on all vs 1% data |  |
| --- | --- | --- | --- | --- | --- |
|  |  | No prior | With prior | No prior | With prior |
| Binary | SPI1 | 95.6 ± 1.0 | **134.3 ± 1.0** | 29.8 ± 0.7 | **42.0 ± 0.8** |
| | GATA2 | 105.8 ± 1.5 | **140.5 ± 1.8** | 29.6 ± 1.3 | **43.5 ± 1.4** |
| | Nanog/Oct4/Sox2 | 107.0 ± 2.5 | **129.0 ± 2.2** | 43.1 ± 1.3 | **53.1 ± 1.7** |
| | K562 | 121.1 ± 2.0 | **197.5 ± 2.5** | 48.3 ± 1.7 | **67.2 ± 1.8** |
| Profile | SPI1 | 332.9 ± 11.2 | **405.6 ± 11.4** | 77.6 ± 4.4 | **233.1 ± 9.8** |
| | GATA2 | 369.1 ± 10.1 | **390.5 ± 9.5** | 102.9 ± 5.2 | **121.6 ± 5.9** |
| | Nanog/Oct4/Sox2 | 216.7 ± 5.7 | **246.5 ± 4.8** | 66.9 ± 4.5 | **109.7 ± 6.2** |
| | K562 | 210.2 ± 5.8 | **236.0 ± 4.2** | 31.4 ± 1.5 | **121.3 ± 5.2** |

## 7 Fourier-based priors increase specificity of model reliance on regions of regulatory function

When we train our models with the Fourier-based prior, we generally observe that the models are able to place higher attribution in high-confidence regions of regulatory significance, and less attribution on other irrelevant areas (Table 3). To show this, we consider test-set peak sequences, and compute the Spearman correlation between each base pair’s DeepSHAP importance to its distance from the peak summit. A more negative correlation implies that the model is placing higher importance closer to the most confident region of binding, as determined by the experimental assay. Additionally, we compute a rank-based measure of specificity by calculating the precision and recall of individual bases in the input sequences that overlap the precise *∼* 200 bp peak regions when ranked by importance (Supplementary Figure S17). A higher area under the precision–recall curve implies that the model places higher importance in called peaks, and in a more specific/precise manner. Using a Wilcoxon rank test, all improvements in summit distance correlation when training with the Fourier-based prior are significant at the 1 × 10^−6^ level.

**Table 3:** Placement of importance on summit/peak regions in test set

|  |  | Correlation of importance to distance from summit |  | auPRC of overlap with called peaks |  |
| --- | --- | --- | --- | --- | --- |
|  |  | No prior | With prior | No prior | With prior |
| Binary | SPI1 | $-0.433 \pm 0.002$ | **$-0.442 \pm 0.002$** | 0.595 | **0.616** |
| | GATA2 | $-0.392 \pm 0.003$ | **$-0.419 \pm 0.003$** | 0.608 | **0.617** |
| | Nanog/Oct4/Sox2 | $-0.509 \pm 0.003$ | **$-0.544 \pm 0.003$** | 0.674 | 0.669 |
| | K562 | $-0.335 \pm 0.005$ | **$-0.360 \pm 0.005$** | 0.587 | **0.594** |
| Profile | SPI1 | $-0.464 \pm 0.005$ | **$-0.490 \pm 0.004$** | 0.398 | **0.470** |
| | GATA2 | $-0.391 \pm 0.006$ | **$-0.437 \pm 0.005$** | 0.430 | **0.497** |
| | Nanog/Oct4/Sox2 | $-0.454 \pm 0.005$ | **$-0.476 \pm 0.005$** | 0.464 | **0.516** |
| | K562 | $-0.392 \pm 0.006$ | **$-0.411 \pm 0.006$** | 0.530 | **0.548** |

For the Nanog/Oct4/Sox2 TF binding and the K562 chromatin accessibility datasets, we use additional independent binding data to further assess the ability of a model to place importance in biologically relevant regions. For Nanog/Oct4/Sox2 binding, we consider high-resolution binding peaks called from independently collected ChIP-nexus experiments [4], and for K562 chromatin accessibility, we consider a set of high-resolution binding footprints explicitly derived from several DNase-seq profiles using independent signal processing methods [12]. These ChIP-nexus peaks and footprints define high-confidence, high-resolution regions where bound regulatory motifs are expected to be found. For these datasets, we further demonstrate that models trained with the Fourier-based prior are able to place importance more specifically in relevant regions, by quantifying the fraction of attributions that overlie a ChIP-nexus peak or DNase-seq footprint, and by considering the precision– recall of important regions overlapping with the ChIP-nexus peaks or DNase-seq footprints (Table 4, Supplementary Figure S18–S19). Using a Wilcoxon rank test, all improvements in importance overlap fraction with ChIP-nexus peaks or DNase-seq footprints when training with the Fourier-based prior are significant at the 1 × 10^−6^ level.

**Table 4:**
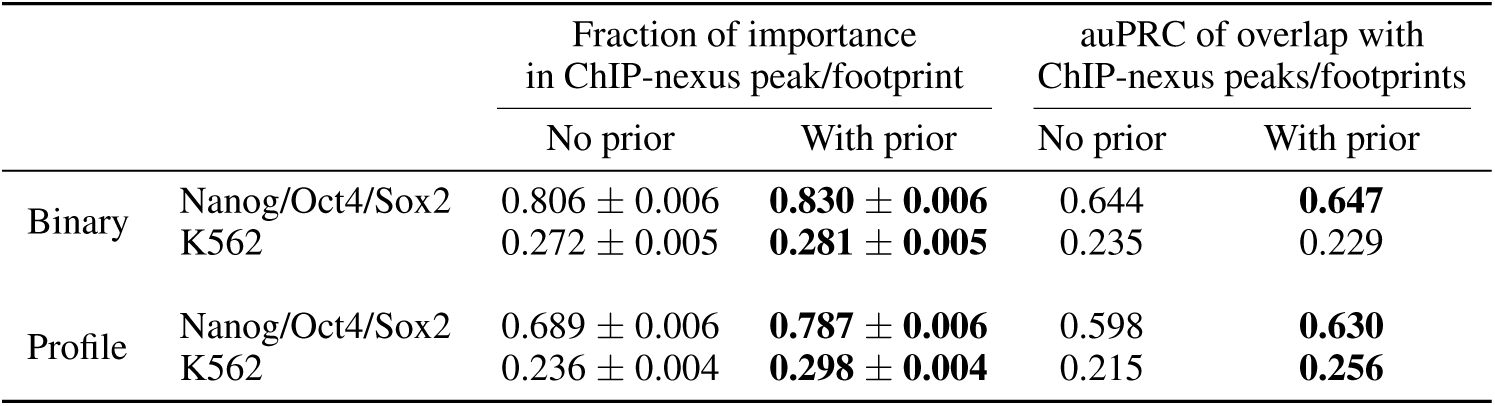
Importance placement on independently derived ChIP-nexus peaks or DNase-seq footprints

We note that the only conditions where the auPRC of overlap with peaks or footprints did not show an improvement when training with the Fourier-based prior were binary models trained to predict Nanog/Oct4/Sox2 binding or K562 chromatin accessibility. These datasets are expected to require more complex motif sequence syntax. Unlike profile models, binary models by nature do not have the resolution in the labels to distinguish directly bound primary motifs from secondary motifs that indirectly contribute to binding and accessibility [13, 14]. For these tasks, the Fourier-based prior elevates the attribution scores of *all* motifs that are predictive of TF binding or accessibility, even when those motifs are not the primary motif underlying a peak or footprint. In concordance with this observation, we find that for these complex binary tasks, the regions outside ChIP-nexus peaks or K562 footprints that are highlighted by the Fourier-based prior contain motifs of important secondary TFs (Supplementary Figure S20) [4].

Finally, we examine models trained on simulated sequences and evaluate the ability of the Fourier-based prior to focus importance on specific individual motifs. On simple binary models trained on simulated binding of the SPI1 TF from synthetic sequences, training with the Fourier-based prior resulted in a significantly higher fraction of importance being placed in motif instances on average (0.659 ± 0.027 vs. 0.218 ± 0.010, *p* < 1 × 10^−6^ by Wilcoxon test). Additionally, models trained with the Fourier-based prior had a much better auPRC of importance-ranked base overlap with motif instances (0.607 vs. 0.422) (Supplementary Figure S22).

## 8 Fourier-based priors improve interpretability beyond standard regularization techniques

Although attribution priors are technically a form of regularization, their mechanism of directly penalizing spurious attributions allows them to improve the interpretability of models beyond what standard regularization techniques can provide. To demonstrate this, we compare SPI1 binary models trained with L2-regularization (i.e. weight decay) versus the Fourier-based prior. We find that DeepSHAP attributions from the model trained with the Fourier-based prior have a significantly lower sum of high-frequency Fourier components (0.377 ± 0.002 vs. 0.400 ± 0.002, *p* < 1 × 10^−6^ by Wilcoxon test) and a significantly lower Shannon entropy (7.891 ± 0.011 vs. 8.908 ± 0.014, *p* < 1 × 10^−6^ by Wilcoxon test) (Supplementary Figure S23). Additionally, the model trained with the Fourier-based prior has a better correlation of base importance to summit distance compared to L2-regularization (− 0.442 ± 0.002 vs. − 0.421 ± 0.002, *p* < 1 × 10^−6^ by Wilcoxon test), and a higher auPRC of ChIP-seq peak overlap on importance-ranked bases (0.616 vs. 0.585) (Supplementary Figure S24)

## 9 Fourier-based priors resist reliance on GC content as a feature

The background GC content (i.e. proportion of G/C bases) of DNA sequences can be an informative feature to differentiate regulatory DNA sequences from other genomic contexts, but reliance on it reduces the interpretability of genomic models, as it obfuscates the underlying motifs driving biological processes. Using simulated DNA sequences with varying amounts of GC bias in the positive training set, we show a simple binary SPI1 binding model trained with the Fourier-based prior resists relying on GC content as a feature, even when surrounding GC content becomes more and more informative (Figure 5).

**Figure 5:**
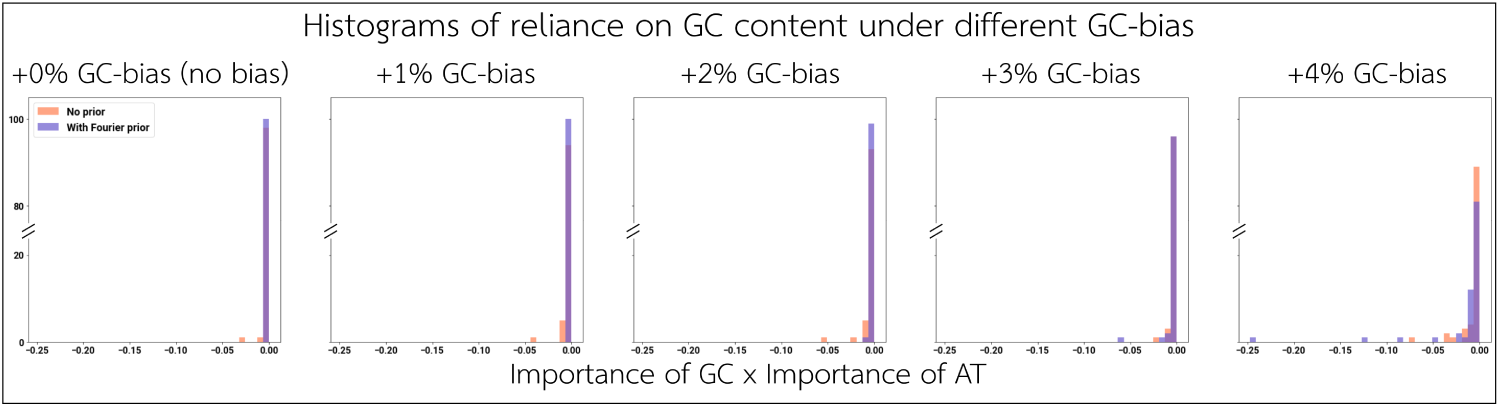
Over different levels of GC bias in the positive training set, we show how heavily models rely on distinguishing G/C and A/T in the background. Background GC content in the positive training set ranges from +0% (the same as the negative set) to an additive +4% more than the negative set. On a set of sampled sequences, the average product of G/C importance and A/T importance in the background sequence reflects how much the model relies on GC content as an informative feature (a more negative product implies a heavier reliance).

Notably, the prior is able to reduce a model’s reliance on informative background GC content, without attenuating the importance of specific GC-rich motifs (Supplementary Figure S25).

## 10 Conclusion

In this work, we introduced and demonstrated the application of a novel attribution prior based on Fourier transforms for improving the stability and interpretability of deep learning models on regulatory DNA. To our knowledge, this is the first application of attribution priors to deep learning models trained on genomic sequence, and the first application of a frequency-based attribution prior in any domain. Our method provides a direct way to train a model with interpretability for scientific discovery as an explicit goal. The Fourier-based prior remains flexibly applied to any architecture, but provides similar gains across models and prediction tasks. Notably, while our prior used input gradients as attributions during training, all the benefits are clearly demonstrated using an alternative reference-based interpretation method (i.e. DeepSHAP).

We showed that models trained with the Fourier-based prior have attributions with a significantly better signal-to-noise ratio, focusing importance primarily on biologically relevant motifs supported by independent data instead of irrelevant background sequence. Hence, models trained with the prior exhibit improved motif discovery and yield motif instance calls that are more likely to underpin regulatory function. Additionally, the Fourier-based prior improves the stability of model learning, directing models to consistently rely on regions of regulatory importance rather than irreproducibly learning background noise. Even when the models are trained with less data, they may be interrogated to discover motifs that are more similar to models that are trained with the full dataset.

Future work could explore other promising attribution prior formulations (e.g. penalizing Shannon entropy in the attributions) and harmoniously merge attribution priors like the Fourier-based prior with more standard forms of regularization. Advances in attribution priors will continue to improve the interpretability of deep learning models in genomics.

## Supplementary Figures and Tables

**Figure S1:**
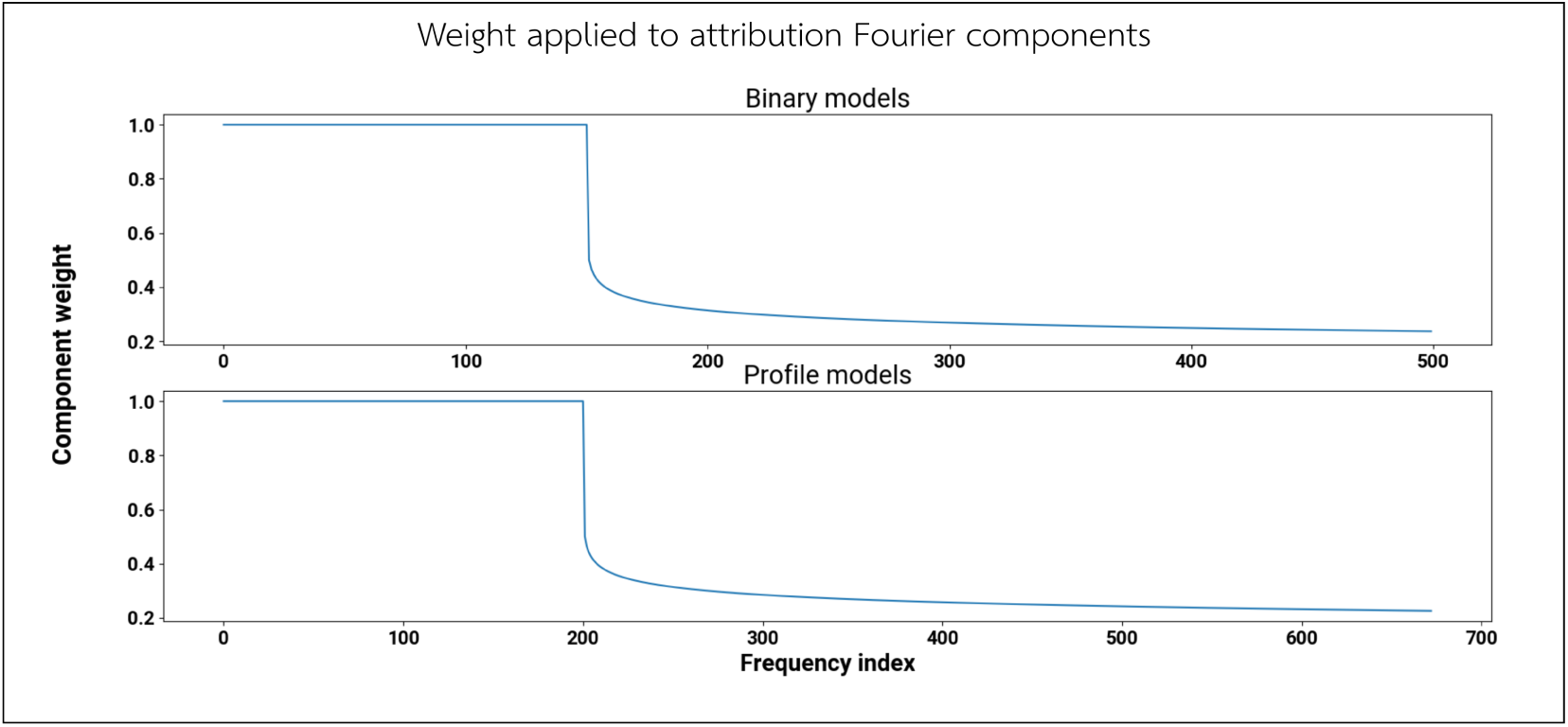
Weight applied to Fourier components in attribution prior loss. To compute the Fourier-based attribution prior loss, the Fourier components corresponding to positive frequencies of the attributions are weighted as shown. This weighted sum constitutes the score of the attributions, and 1 minus this score becomes the attribution prior loss value. Note that because input sequences to binary and profile models have different lengths, the discrete Fourier transform components also have different lengths; in both cases, the frequency threshold *T* corresponds to a minimum expected motif length of 6–7.

**Figure S2:**
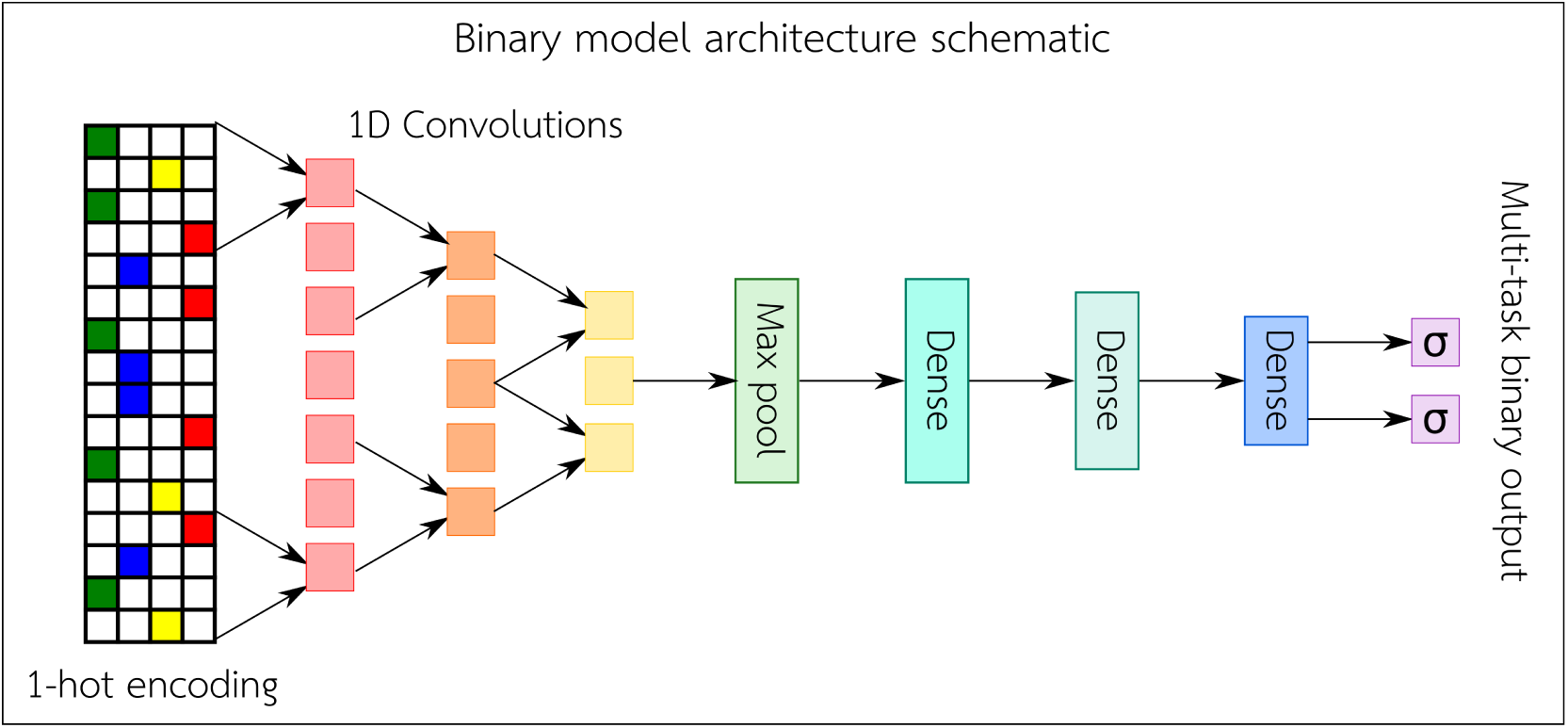
Schematic of binary model architecture. A one-hot encoded sequence is fed into three consecutive convolutional layers. The resulting activations are passed through a max pooling layer, followed by three dense layers, where the final dense layer outputs a sigmoid-transformed binary prediction.

**Figure S3:**
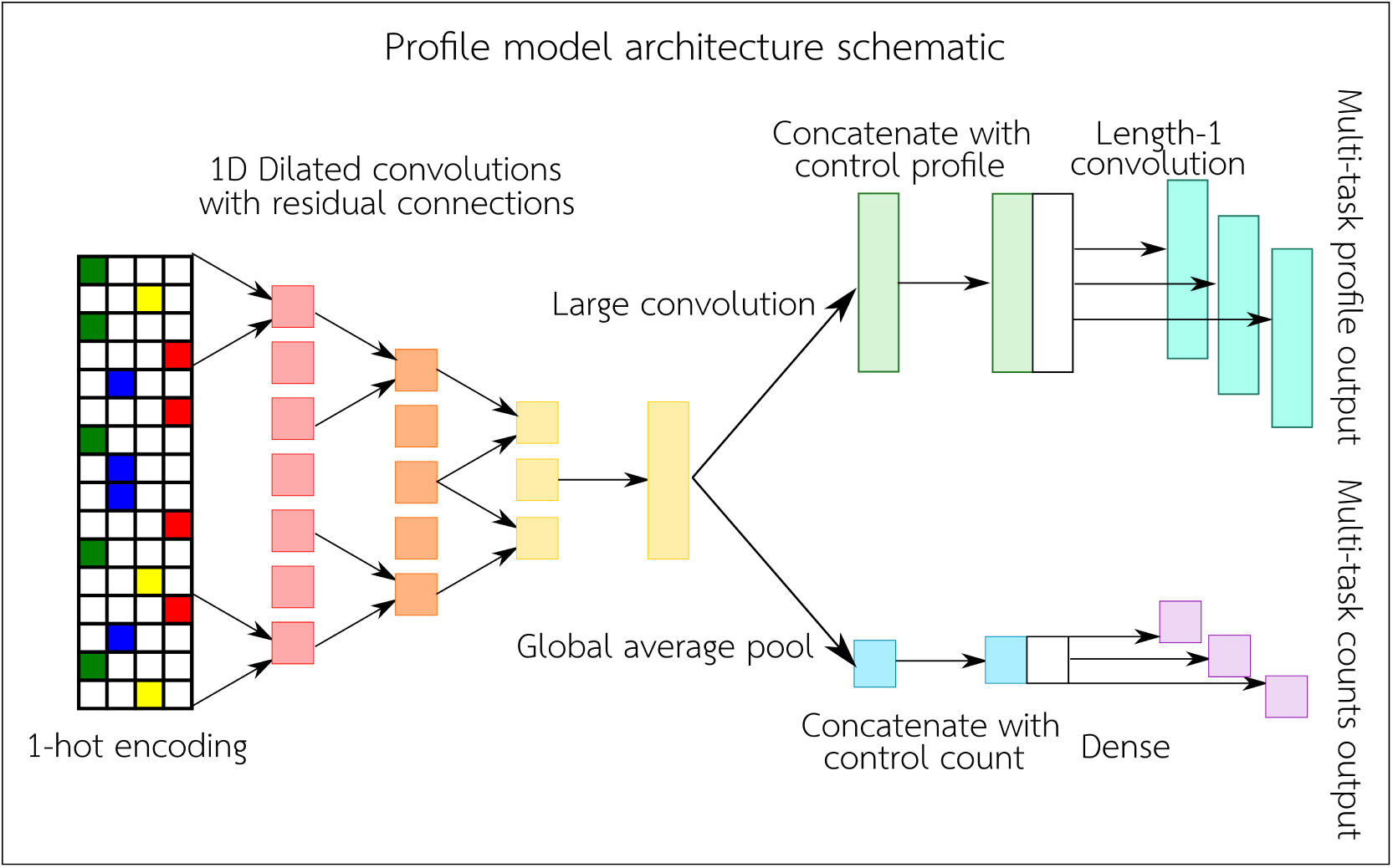
Schematic of profile model architecture, based on the architecture in Avsec et al. [4]. A one-hot encoded sequence is fed into six consecutive dilated convolutional layers with summed residual connections. For each task, the model predicts a profile shape and a read count. The profile shape prediction is obtained by feeding the activations from the dilated convolutions to another convolutional layer with a large kernel size, concatenating the result with a set of control profiles, and performing a length-1 convolution over the concatenation to yield a profile shape prediction. The read count prediction is obtained by feeding the activations from the dilated convolutions through a global average pooling layer, concatenating the result with a set of control read counts, and passing this concatenation through a dense layer to obtain predicted read counts.

**Figure S4:**
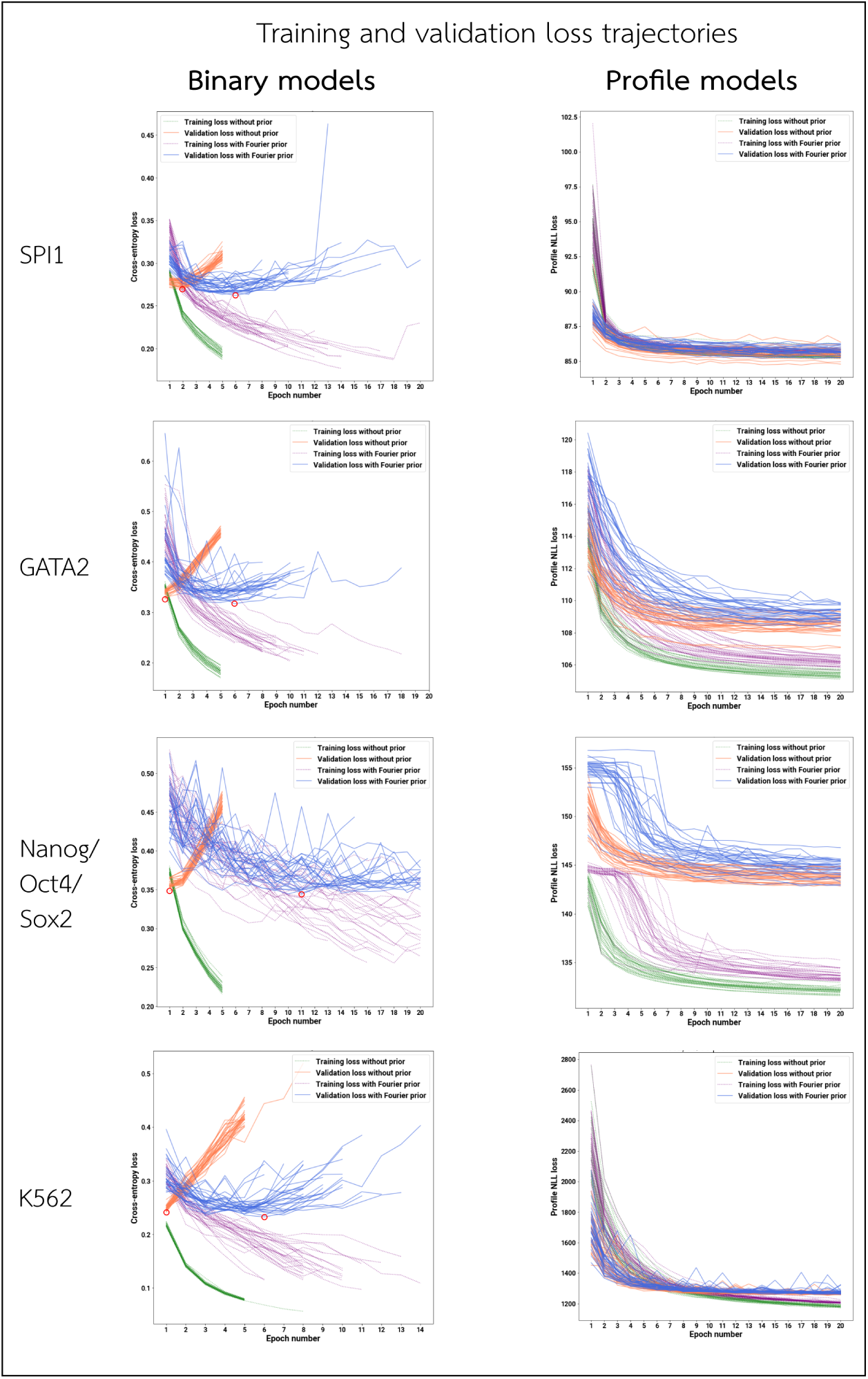
Training and validation correctness loss trajectories. For each architecture and dataset, we show the trajectory of the training and validation correctness losses (i.e. excluding any attribution prior loss) after each epoch of training, over all random initializations. Loss values shown begin after the first epoch of training. In general, binary models (left) overfit very easily, with validation loss visibly growing after the first epoch. Profile models (right), however, are much more resilient to overfitting, as they benefit from extensive data augmentation through random jitters in the input sequences. On binary models (left), red circles indicate the models that achieve the lowest validation loss, trained with and without the Fourier-based prior.

**Table S1:**
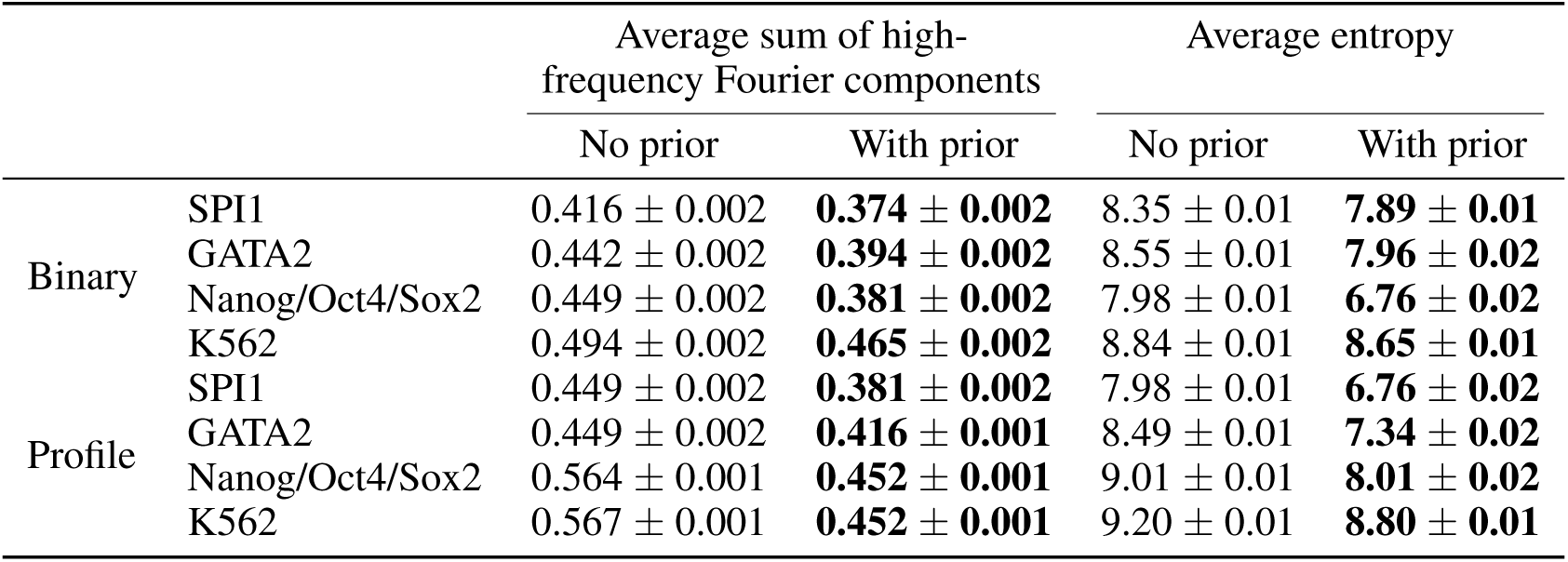
Improved interpretability on test-set sequences

|  |  | Average sum of high-frequency Fourier components |  | Average entropy |  |
| --- | --- | --- | --- | --- | --- |
|  |  | No prior | With prior | No prior | With prior |
| Binary | SPI1 | $0.416 \pm 0.002$ | **$0.374 \pm 0.002$** | $8.35 \pm 0.01$ | **$7.89 \pm 0.01$** |
| | GATA2 | $0.442 \pm 0.002$ | **$0.394 \pm 0.002$** | $8.55 \pm 0.01$ | **$7.96 \pm 0.02$** |
| | Nanog/Oct4/Sox2 | $0.449 \pm 0.002$ | **$0.381 \pm 0.002$** | $7.98 \pm 0.01$ | **$6.76 \pm 0.02$** |
| | K562 | $0.494 \pm 0.002$ | **$0.465 \pm 0.002$** | $8.84 \pm 0.01$ | **$8.65 \pm 0.01$** |
| Profile | SPI1 | $0.449 \pm 0.002$ | **$0.381 \pm 0.002$** | $7.98 \pm 0.01$ | **$6.76 \pm 0.02$** |
| | GATA2 | $0.449 \pm 0.002$ | **$0.416 \pm 0.001$** | $8.49 \pm 0.01$ | **$7.34 \pm 0.02$** |
| | Nanog/Oct4/Sox2 | $0.564 \pm 0.001$ | **$0.452 \pm 0.001$** | $9.01 \pm 0.01$ | **$8.01 \pm 0.02$** |
| | K562 | $0.567 \pm 0.001$ | **$0.452 \pm 0.001$** | $9.20 \pm 0.01$ | **$8.80 \pm 0.01$** |

**Figure S5:**
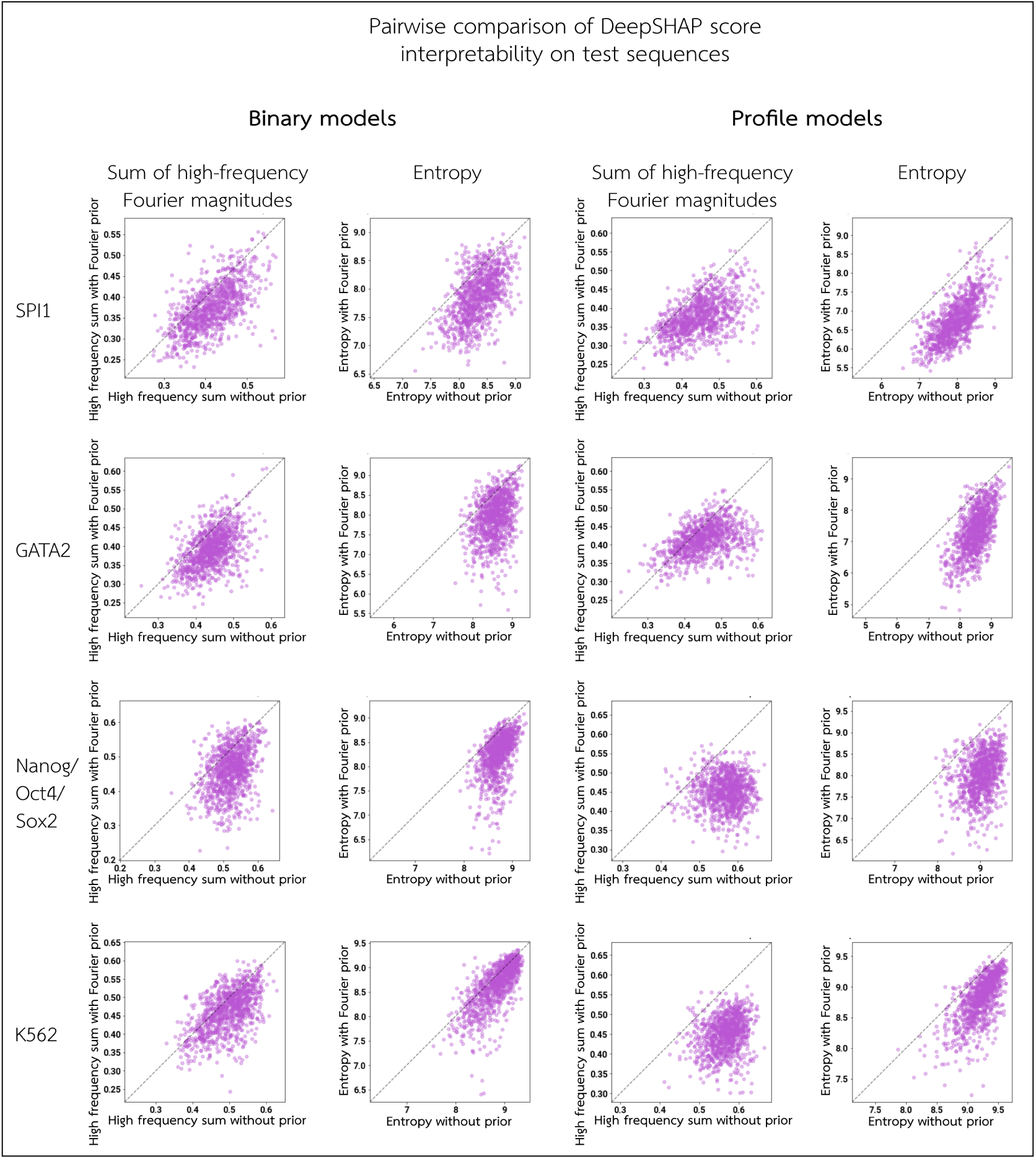
Signal-to-noise ratio of attributions across test set sequences. For each architecture and dataset, we compute DeepSHAP importance scores for 1000 randomly selected peak sequences from the test set. An improvement in the signal-to-noise ratio of the attributions is quantified as a reduction in the high-frequency Fourier component magnitudes, and as a reduction in Shannon entropy. We compare the sum of normalized high-frequency Fourier components and the Shannon entropy for each sequence, between models trained with versus without the Fourier-based prior.

**Figure S6:**
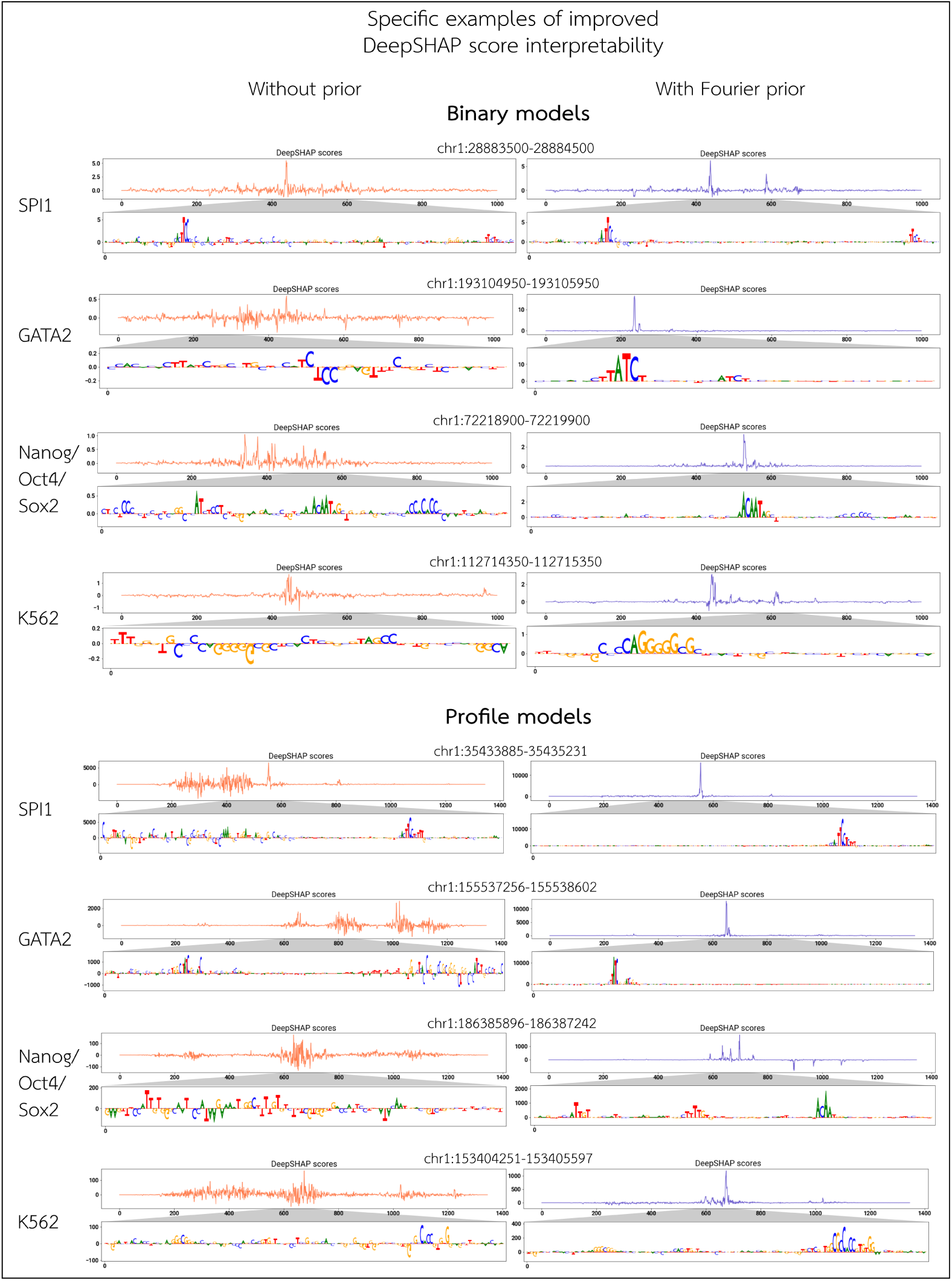
Specific examples of improved signal-to-noise ratio on DeepSHAP scores. For each architecture and dataset, we show the DeepSHAP attribution scores of specific peak sequences. For each selected input sequence, we display the value of the DeepSHAP importance for the bases present along the entire input sequence, as well as the base-pair-level attributions in the summit region.

**Figure S7:**
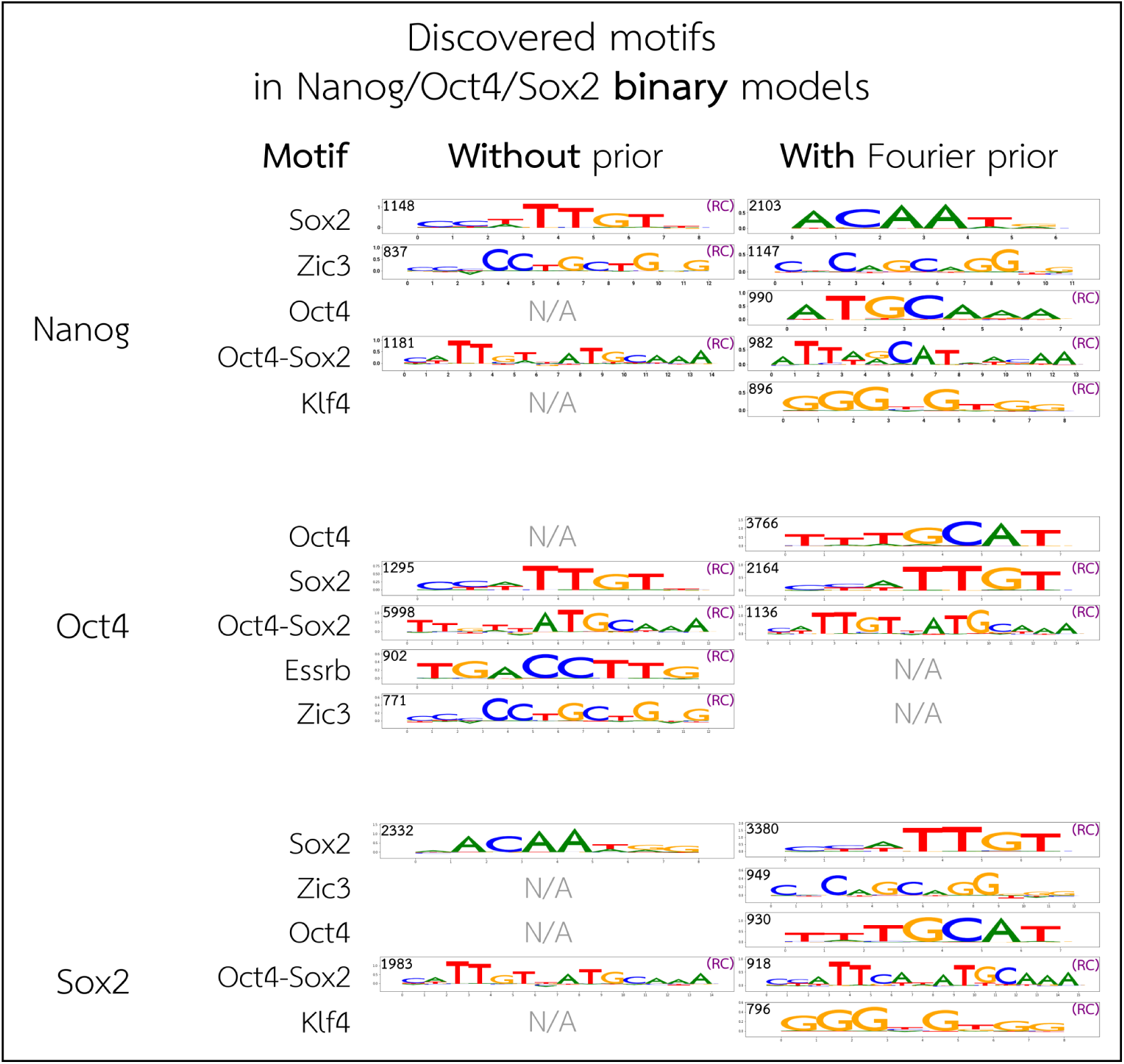
Discovered motifs from Nanog/Oct4/Sox2 binary models. For each TF, we utilize TF-MoDISco to discover motifs using DeepSHAP attributions of test-set peak sequences. We show the relevant motifs identified by TF-MoDISco which pass our thresholds (Supplementary Methods Sec. 5), and match them to known motifs from Avsec et al. [4]. “(RC)” denotes that the motif shown is reverse-complemented relative to the orientation in Avsec et al. [4]. The number in the top left of each motif indicates the number of seqlets identified by TF-MoDISco that underlie the motif.

**Figure S8:**
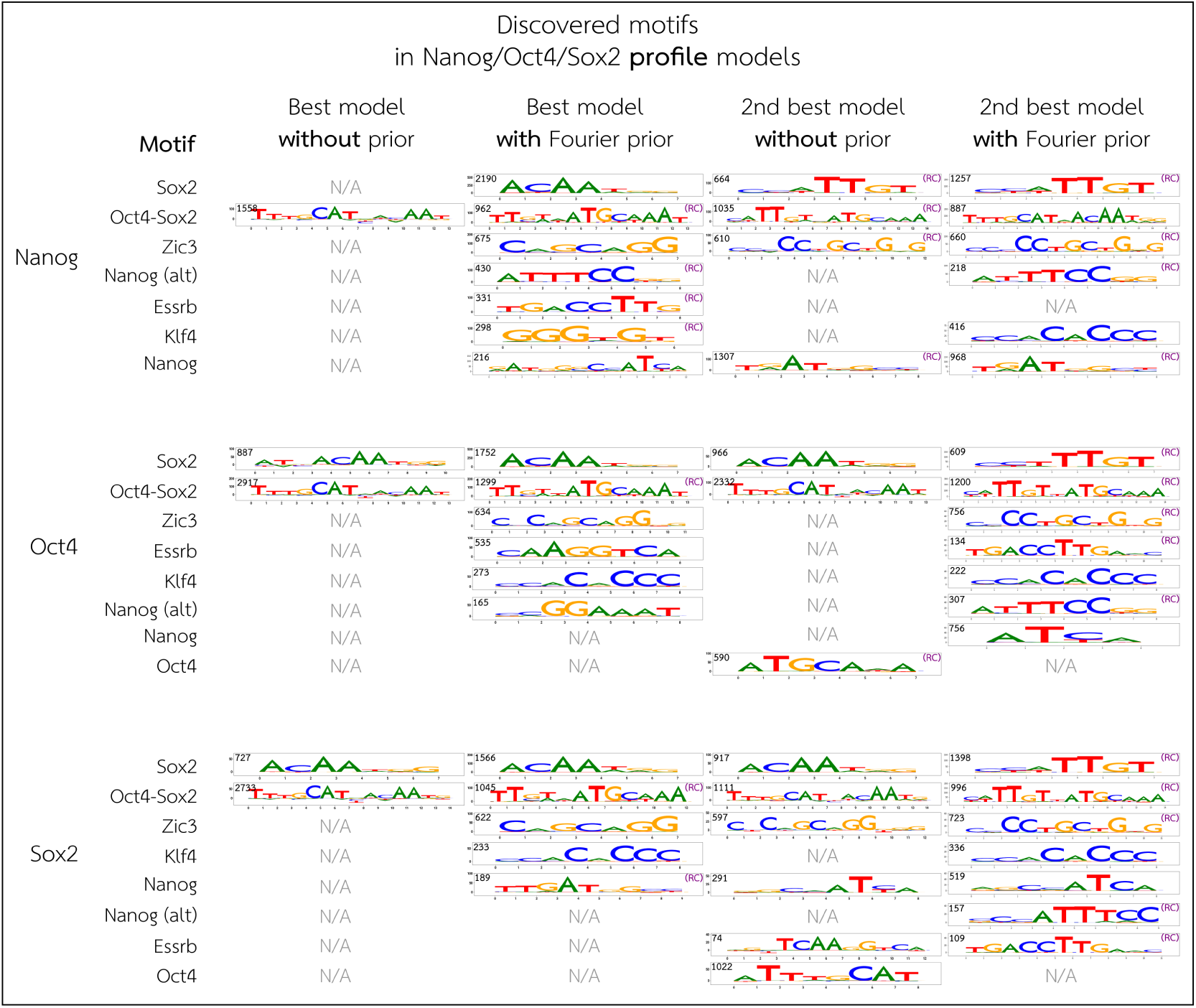
Discovered motifs from Nanog/Oct4/Sox2 profile models. For each TF, we utilize TF-MoDISco to discover motifs using DeepSHAP attributions of test-set peak sequences. We show the relevant motifs identified by TF-MoDISco which pass our thresholds (Supplementary Methods Sec. 5), and match them to known motifs from Avsec et al. [4]. “(RC)” denotes that the motif shown is reverse-complemented relative to the orientation in Avsec et al. [4]. The number in the top left of each motif indicates the number of seqlets identified by TF-MoDISco that underlie the motif. While we focus our downstream analysis on the best-performing models with and without the Fourier-based prior, the best-performing profile model trained without the prior identified much fewer motifs than the model trained with the prior, so we also show the motifs identified using TF-MoDISco on the second-best-performing models, both with and without the prior. This demonstrates that the profile models without the Fourier-based prior are capable of learning the larger set of motifs expected, but are limited by noisy and irreproducible attributions.

**Figure S9:**
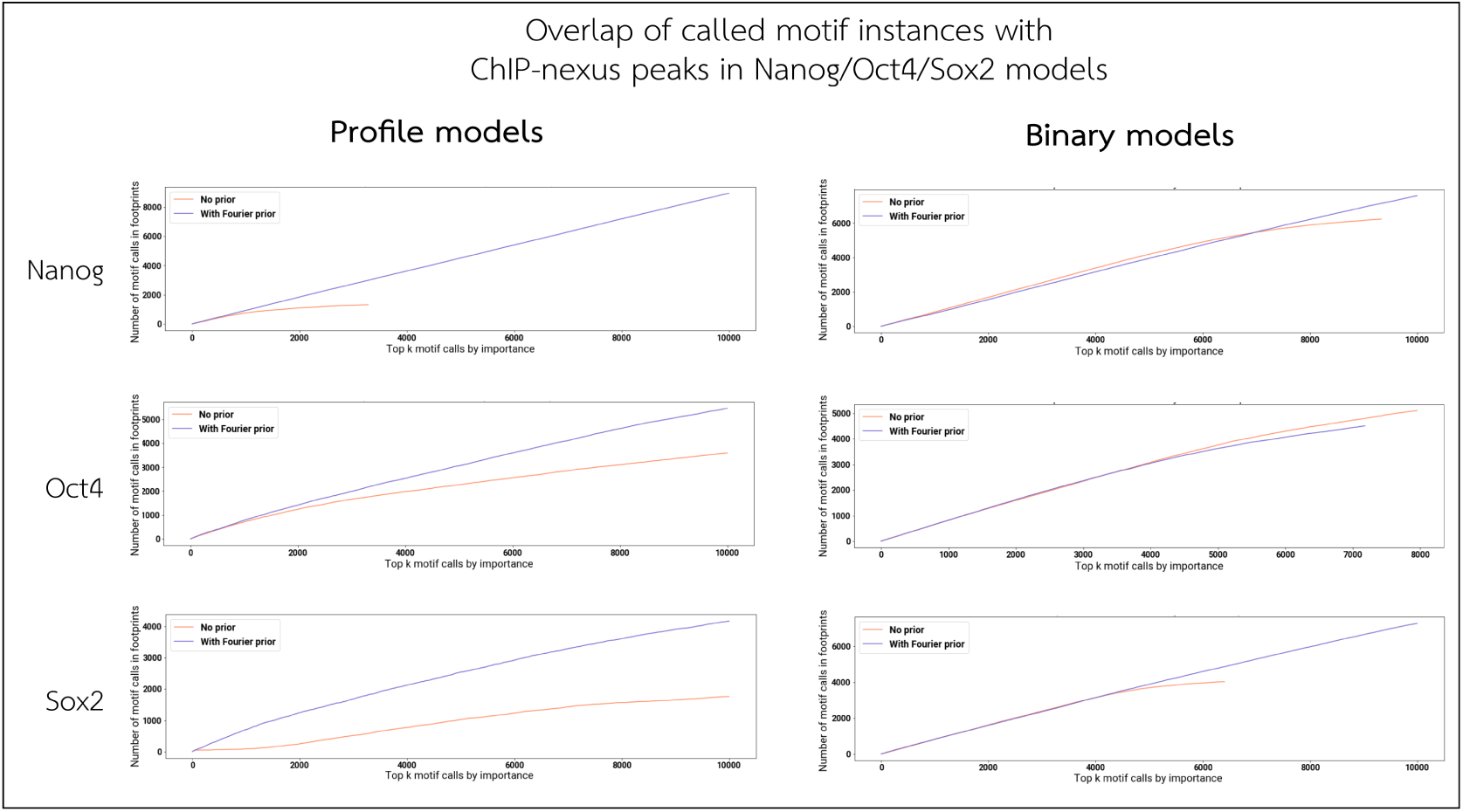
Motif instance call support. For each TF in the Nanog/Oct4/Sox2 models, we perform motif instance calling using the discovered motifs on a sample of 1000 test-set peak sequences. We rank the motif instance calls by total DeepSHAP importance, and compute a cumulative count of how many instances overlap with a ChIP-nexus peak for that TF. Note that the models trained without the prior typically have fewer motif calls in total (due to lower-quality attribution scores and fewer motifs discovered by TF-MoDISco), resulting in the shorter red lines.

**Figure S10:**
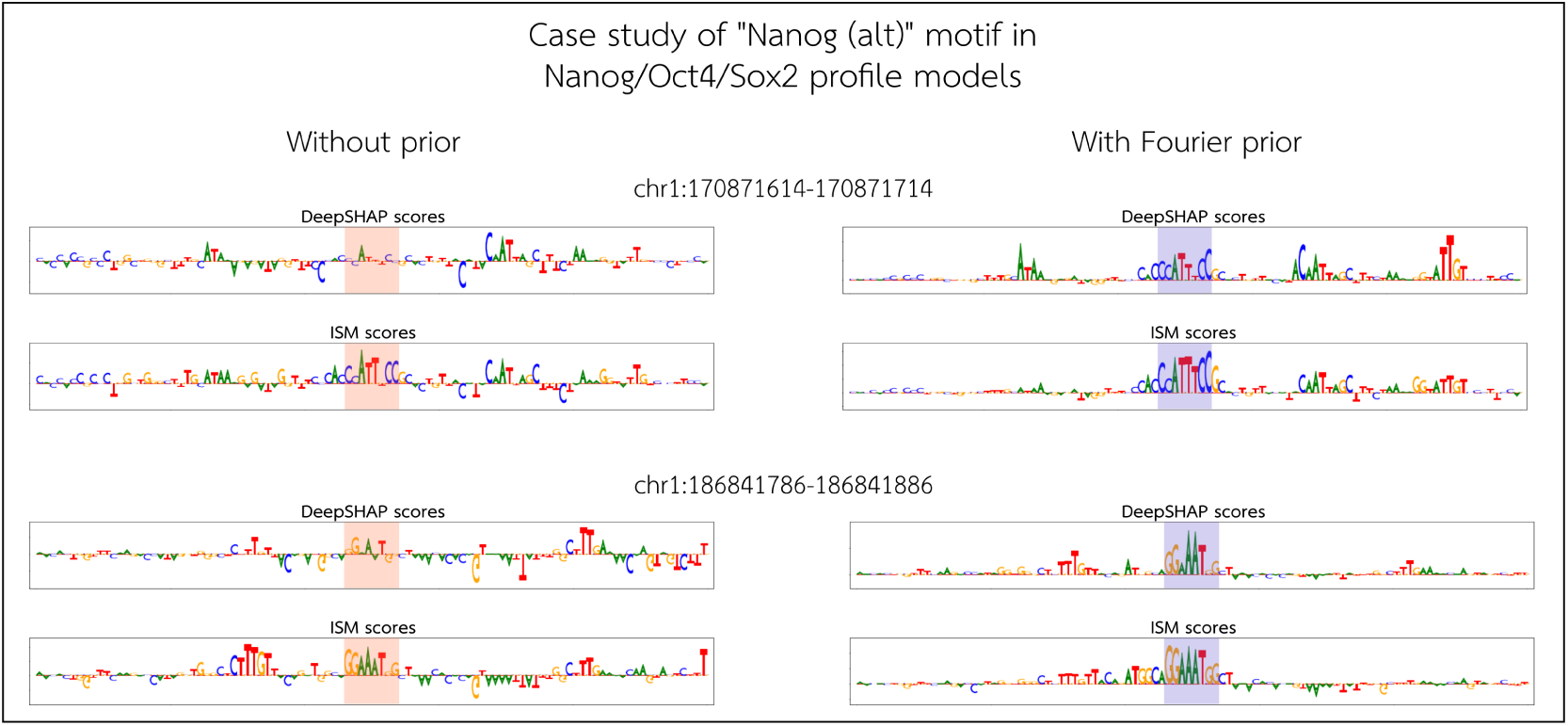
Case study of the “Nanog (alt)” motif in Nanog/Oct4/Sox2 profile models. TF-MoDISco identified the GGAAA “Nanog (alt)” motif from the profile Nanog/Oct4/Sox2 model trained with the Fourier-based prior, but not from the model trained without the prior, even on the Nanog prediction task specifically (Supplementary Figure S8). Focusing on the Nanog prediction task, we show examples of sequences where TF-MoDISco identified a Nanog (alt) motif from the model trained with the prior, and show the importance scores of the same sequence from the model without the prior. We show both the DeepSHAP importance scores, and the perturbation scores derived from *in silico* mutagenesis (ISM). The highlighted regions indicate the location of the Nanog (alt) motif. Notably, ISM scores from the model trained without the prior are generally noisier compared to the model trained with the prior. More importantly, this demonstrates that the model trained without the prior is learning the Nanog (alt) motif, but it is not visible from DeepSHAP importance scores. The model trained with the Fourier-based prior, however, clearly highlights this motif using both methods of interpretation (i.e. DeepSHAP and ISM). This indicates that the Fourier-based prior allows the model to reveal its learned motifs in a human-interpretable way, especially when it is too computationally expensive to rely on perturbation-based scoring methods like ISM, which take orders of magnitude longer to run than DeepSHAP.

**Figure S11:**
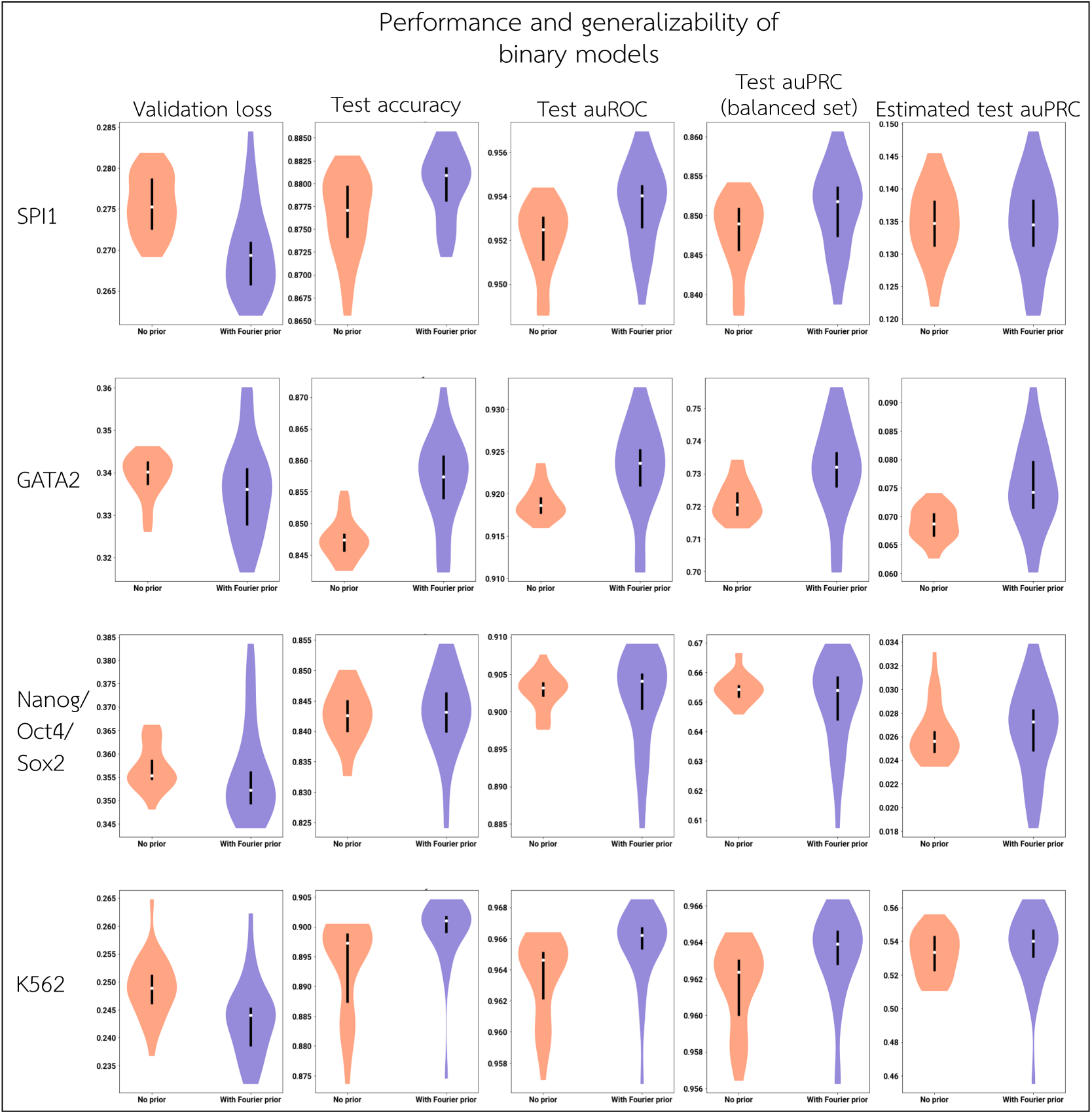
Validation loss and test set performance of binary models. For each dataset, we consider the validation and test set performance of models trained with and without the Fourier-based prior, over 30 random initializations each. Validation loss is computed over all positive examples in the validation set and a equal-sized sample of negative validation examples. Test accuracy, test auROC, and test auPRC are computed over all positive examples in the test set and a equal-sized sample of test negative examples. “Estimated test auPRC” is an estimated measure of auPRC on the full test set without subsampling the negative examples, and is computed by artificially inflating the false positive rate (Supplementary Methods Sec. 2.5).

**Figure S12:**
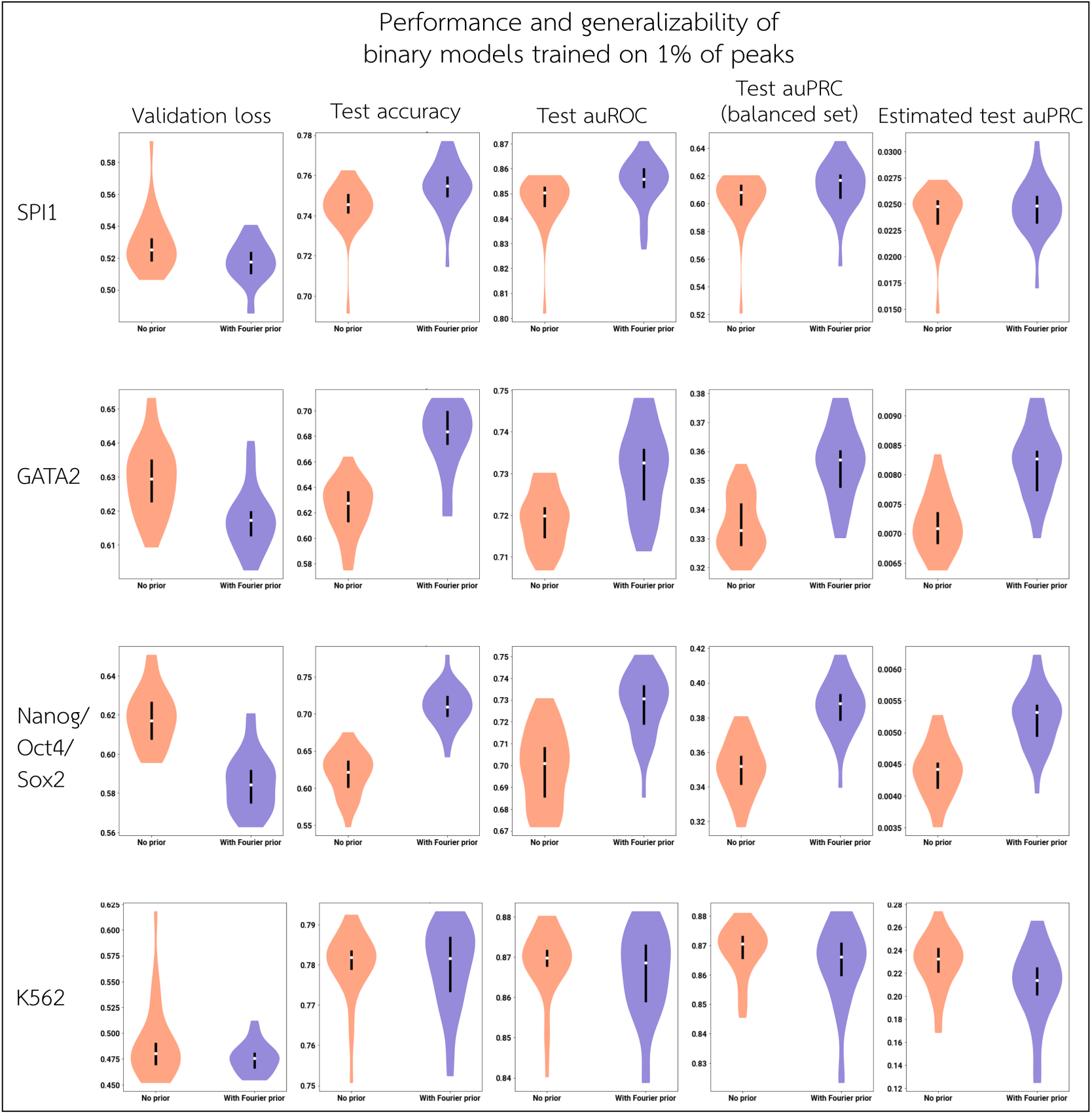
Validation loss and test set performance of binary models on sparse training sets. For each dataset, we consider the validation and test set performance of models trained with and without the Fourier-based prior (on only 1% of the training set), over 30 random initializations each. Validation loss is computed over all positive examples in the validation set and a equal-sized sample of negative validation examples. Test accuracy, test auROC, and test auPRC are computed over all positive examples in the test set and a equal-sized sample of test negative examples. “Estimated test auPRC” is an estimated measure of auPRC on the full test set without subsampling the negative examples, and is computed by artificially inflating the false positive rate (Supplementary Methods Sec. 2.5)

**Figure S13:**
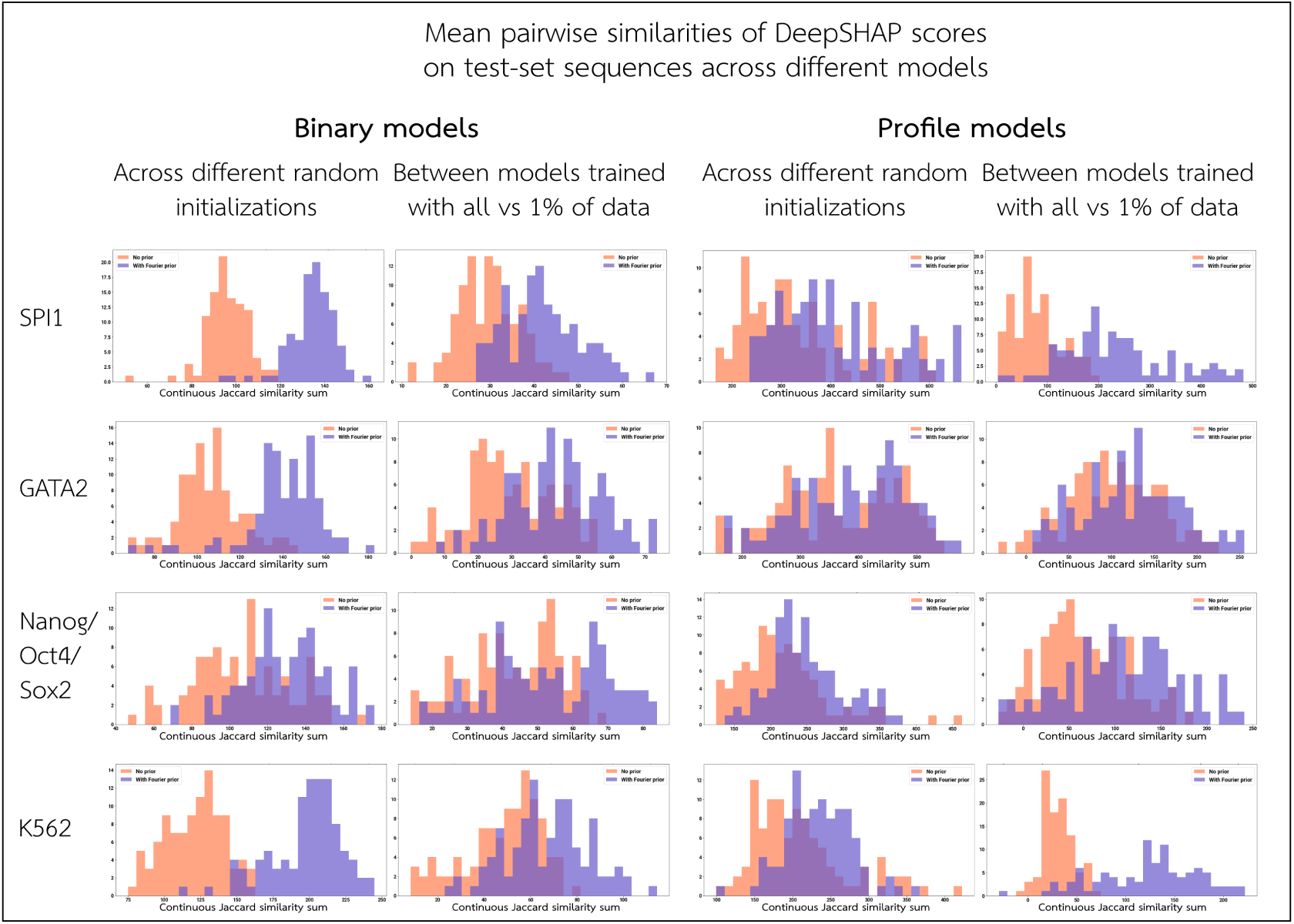
Stability of DeepSHAP scores across different models on test-set sequences. For each architecture and dataset, we sample 100 peak sequences from the test set and compute the DeepSHAP attributions for the sequence for multiple models. For each sequence, we compute the pairwise similarity of the attributions between 30 random initializations (left), or between the top 5 models trained with all of the data versus the top 5 trained with only 1% of the data (right). We quantify attribution similarity by computing the continuous Jaccard score at each base, and summing the scores across the sequence.

**Figure S14:**
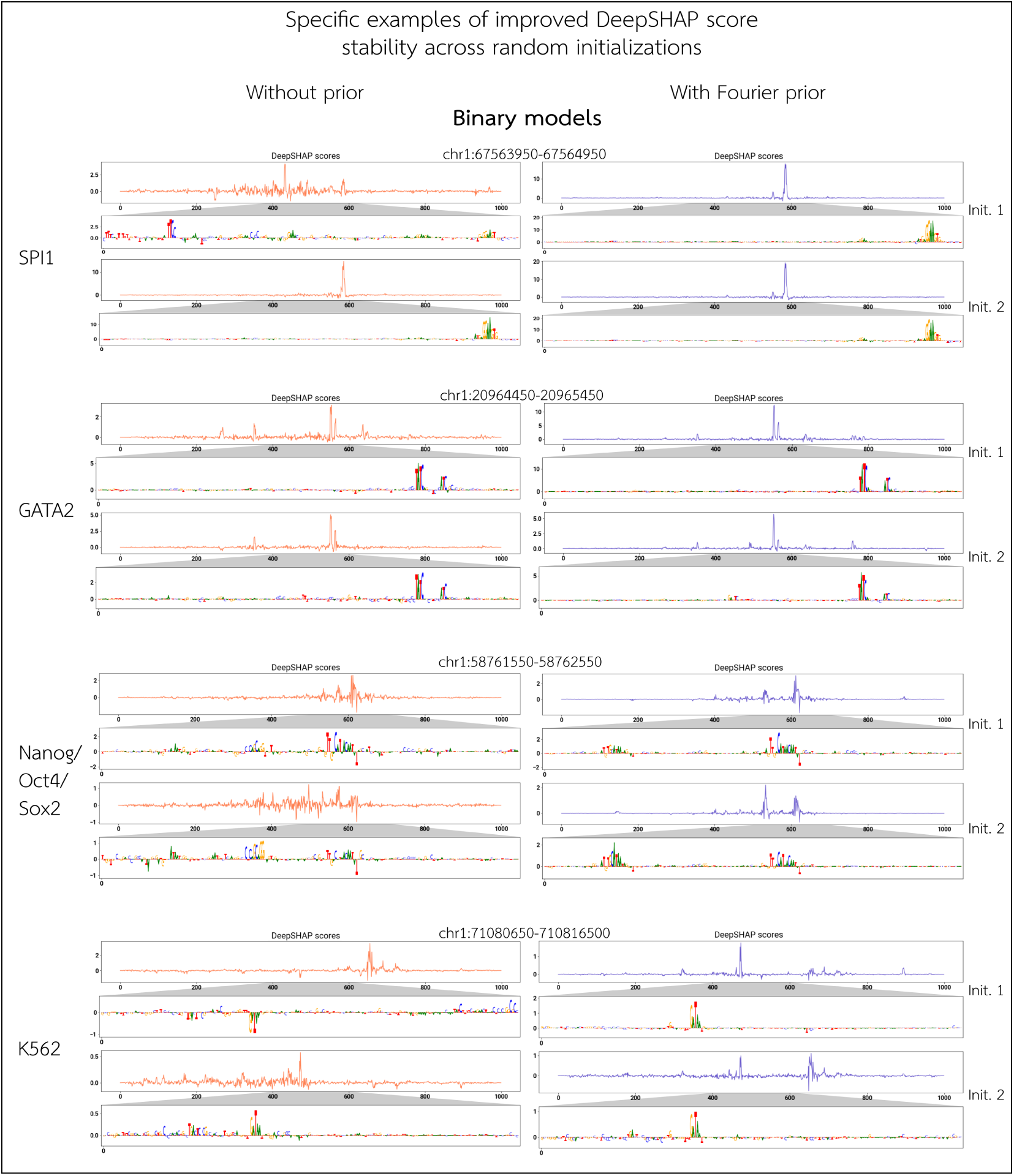
Specific examples of DeepSHAP attribution stability across different random initializations (binary models). For each binary model, we show an example of the DeepSHAP attributions on a test-set sequence between a pair of models of different random initializations, comparing models trained with versus without the Fourier-based prior.

**Figure S15:**
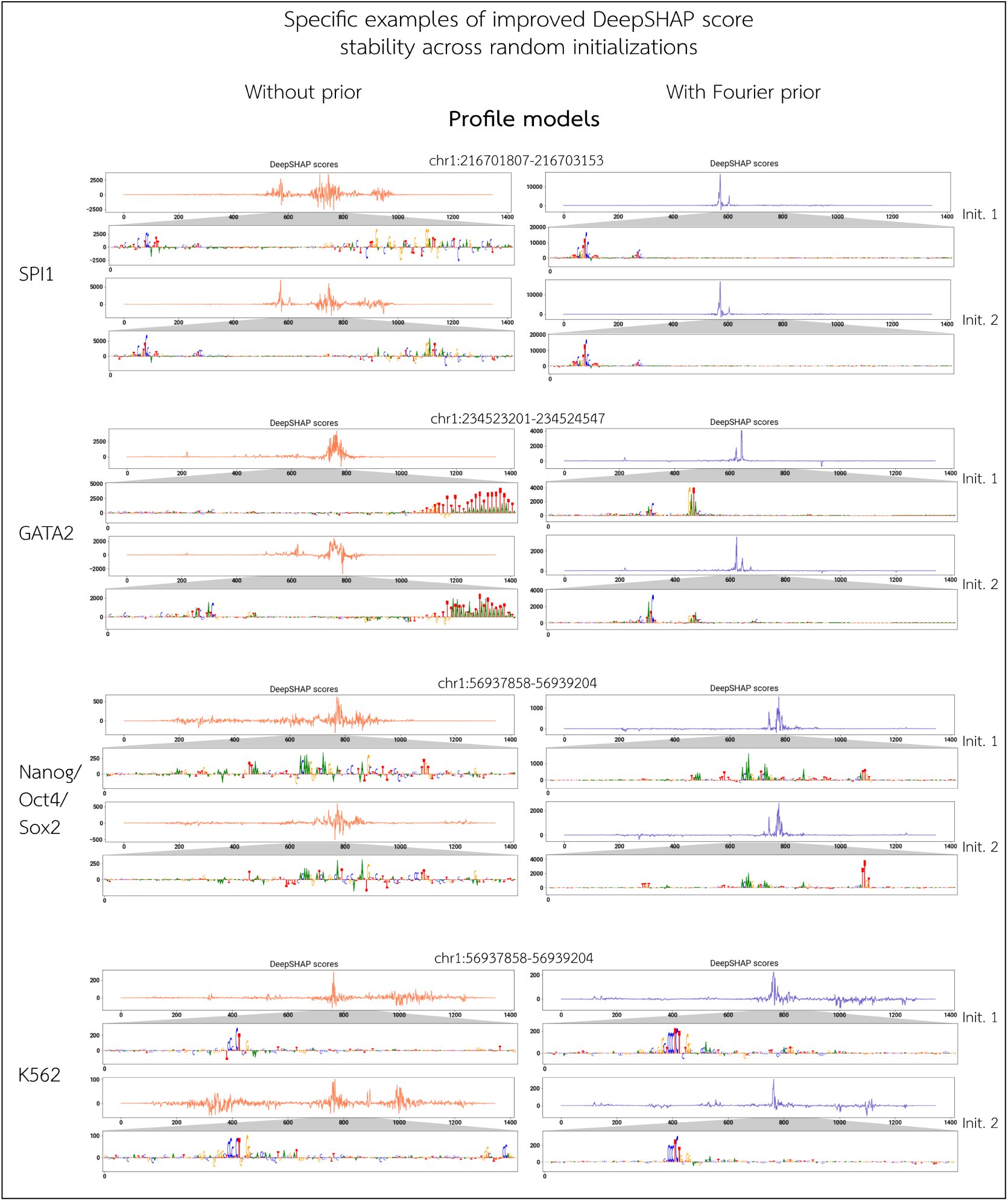
Specific examples of DeepSHAP attribution stability across different random initializations (profile models). For each profile model, we show an example of the DeepSHAP attributions on a test-set sequence between a pair of models of different random initializations, comparing models trained with versus without the Fourier-based prior.

**Figure S16:**
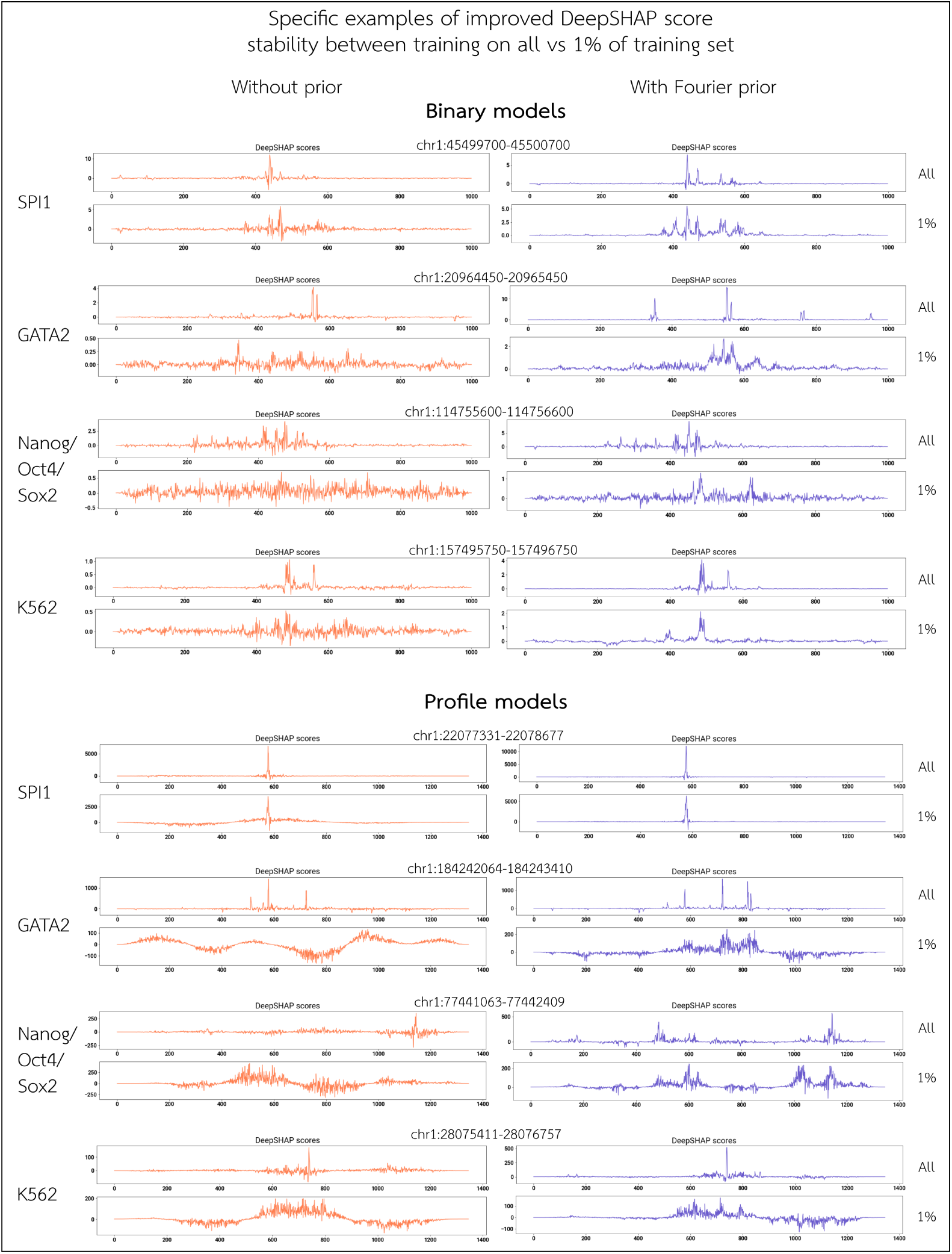
Specific examples of DeepSHAP attribution stability between training on all versus 1% of the training set. For each architecture and dataset, we show an example of the DeepSHAP attributions on a test-set sequence between a model trained on all and only 1% of the training set, comparing models trained with versus without the Fourier-based prior.

**Figure S17:**
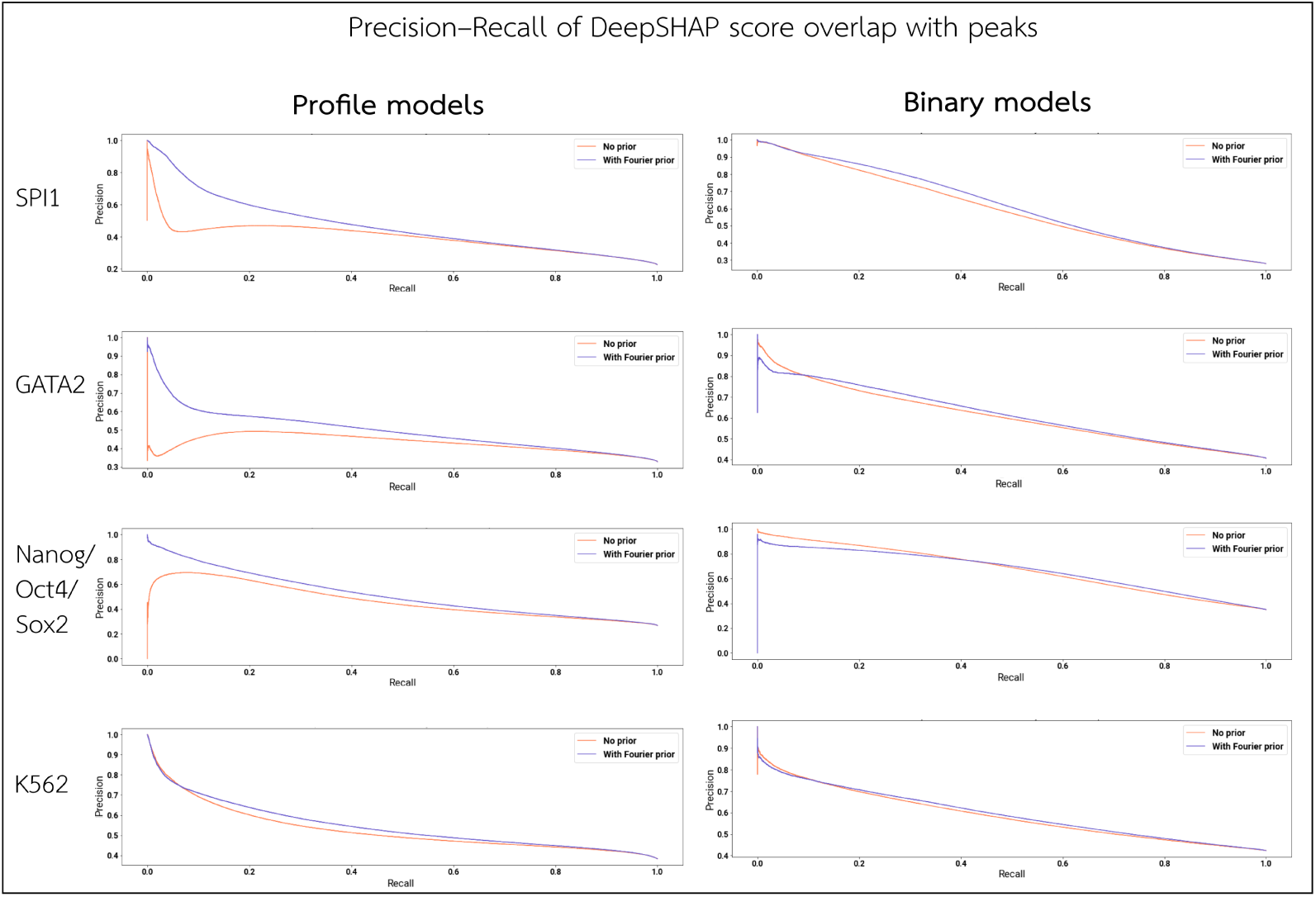
Precision–recall of importance-ranked base overlap with called training peaks. For each architecture and dataset, we sample 1000 peak sequences from the test set and compute the DeepSHAP attributions. We rank bases in descending order of total importance, and generate a precision–recall curve by treating the set of bases that overlap an underlying ChIP-seq or DNase-seq peak as “positives”. See Table 3 in the main text for the corresponding auPRC values.

**Figure S18:**
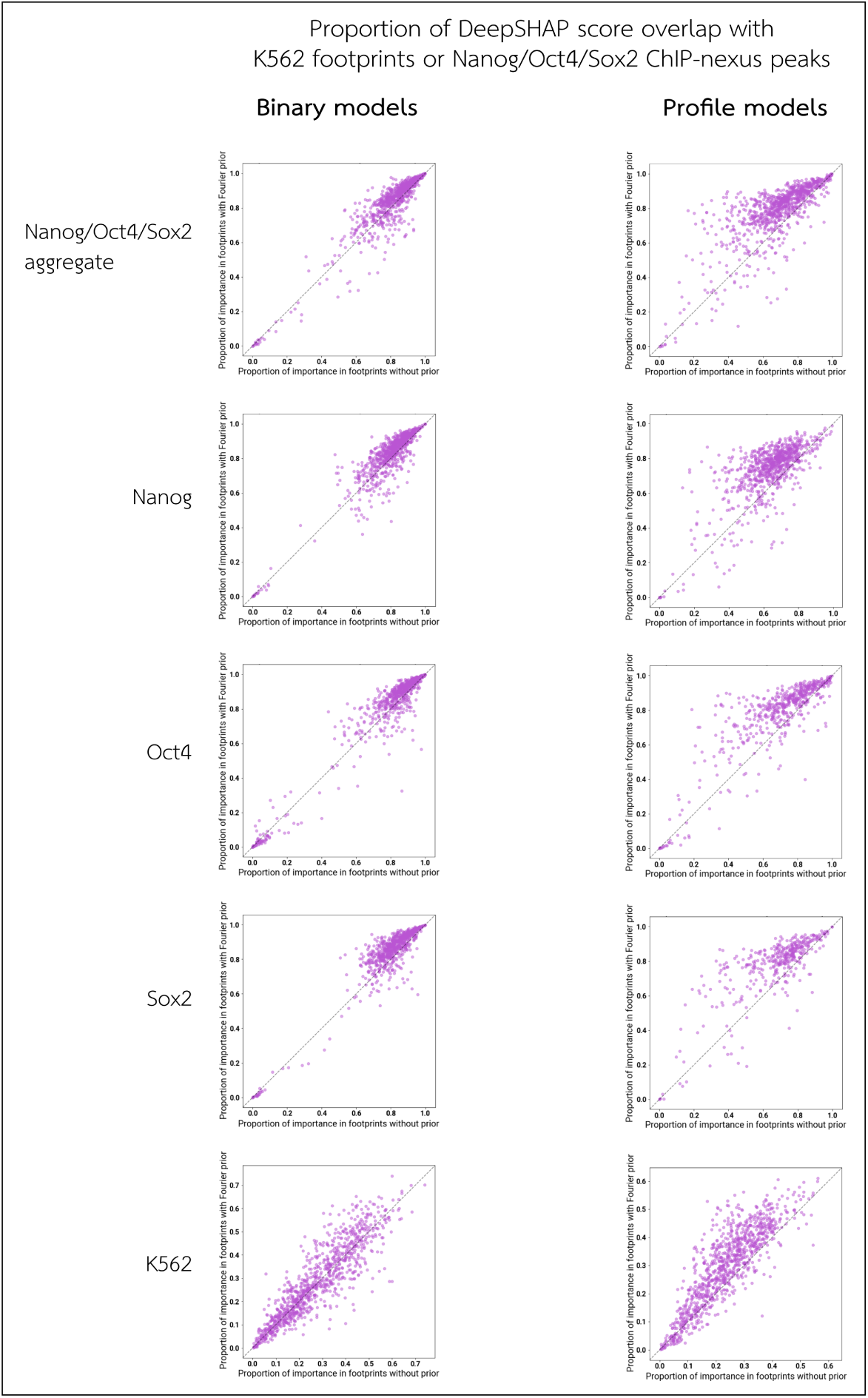
Fraction of importance in ChIP-nexus peaks or DNase footprints. For Nanog/Oct4/Sox2 TF ChIP-seq or K562 DNase-seq models, we sample 1000 peak sequences from the test set and compute the fraction of total DeepSHAP importance that overlaps a Nanog/Oct4/Sox2 ChIP-nexus peak or K562 footprint. For each input sequence, we compare the proportion of attribution by magnitude overlapping a ChIP-nexus peak or footprint when a model is trained with versus without the Fourier-based prior. For Nanog/Oct4/Sox2 models, we also show the overlap of task-specific importance on the set of corresponding ChIP-nexus peaks. See Table 4 in the main text for the corresponding average values; in all cases, the models trained with the prior place a significantly higher fraction of total importance in the peaks/footprints.

**Figure S19:**
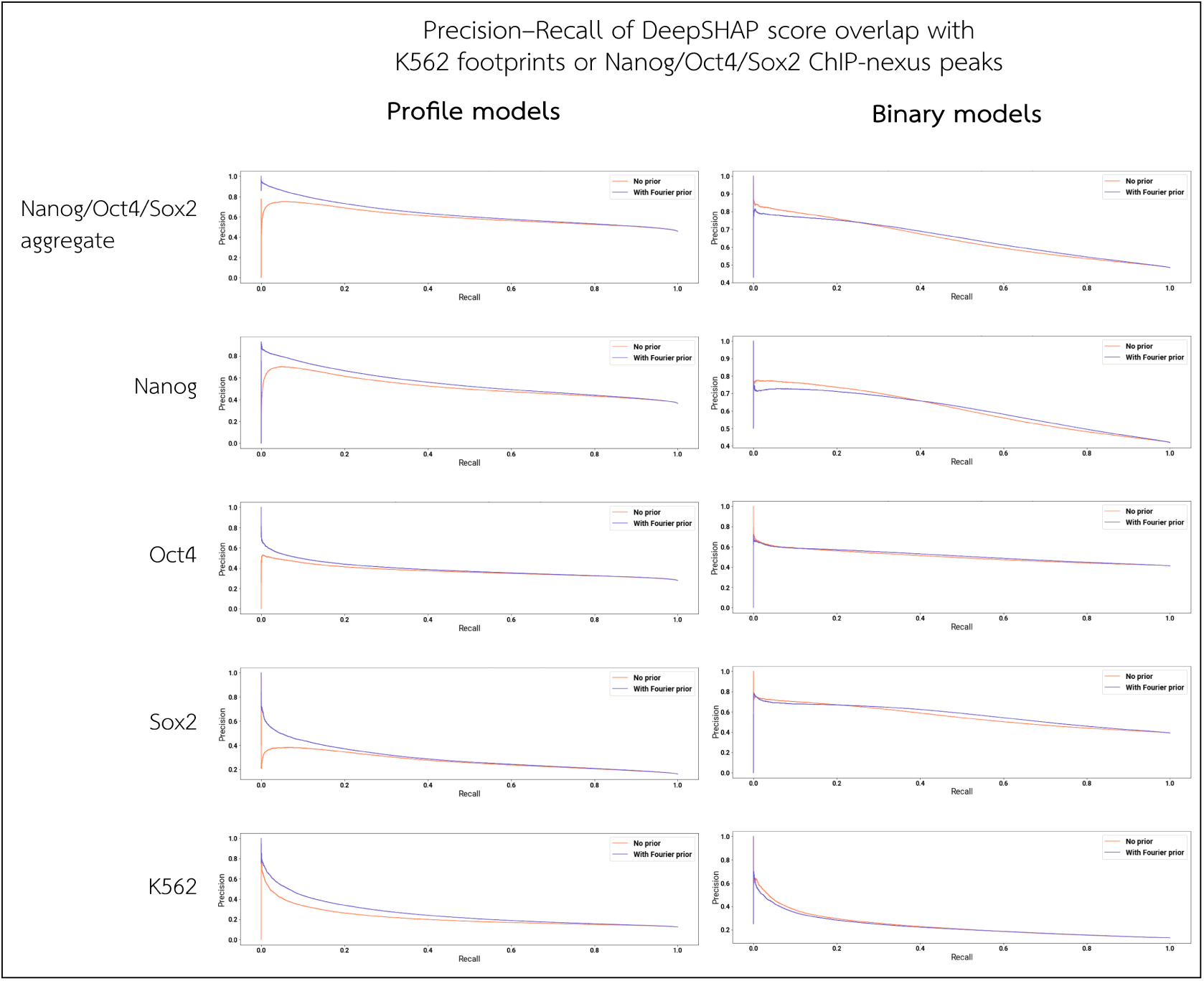
Precision–recall of important bases overlapping with ChIP-nexus peaks or DNAse footprints. For models trained to predict Nanog/Oct4/Sox2 TF ChIP-seq or K562 DNase-seq, we sample 1000 peak sequences from the test set and compute the DeepSHAP importance. We rank bases in descending order of total importance and generate a precision–recall curve by treating the set of bases that overlap a Nanog/Oct4/Sox2 ChIP-nexus peak or K562 DNase-seq footprint as “positives”. For Nanog/Oct4/Sox2 models, we show these curves both using a ranking based on the total importance across all three tasks (in which case we use the union of ChIP-nexus peaks over the three tasks for the labels), as well as a ranking based on the importance scores of each individual task (in which case we use only the ChIP-nexus peaks of the corresponding task for the labels). See Table 4 in the main text for the corresponding auPRC values.

**Figure S20:**
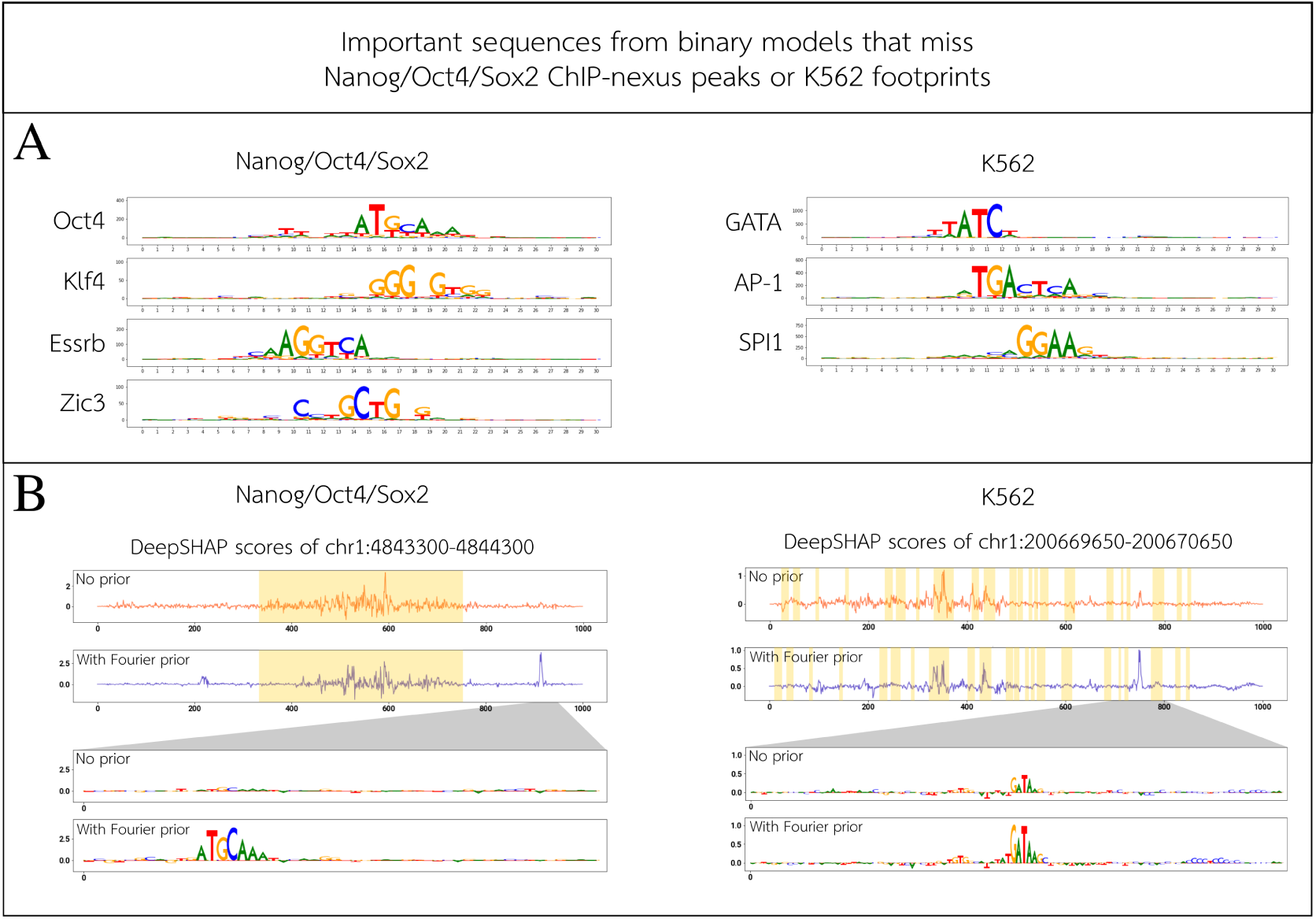
Motifs in important regions outside of peaks. For the Nanog/Oct4/Sox2 TF ChIP-seq and K562 DNase-seq binary models, we examine regions where models trained with the Fourier-based prior place high DeepSHAP importance, yet do not overlap Nanog/Oct4/Sox2 ChIP-nexus peaks or K562 footprints, respectively. **A)** We cluster these regions using the TF-MoDISco clustering algorithm, and show the PWMs of top motif clusters, along with annotations of relevant TFs that are associated to each motif. **B)** We show some specific examples where a model trained with the Fourier-based prior places higher importance (relative to the model trained without the prior) outside of a ChIP-nexus peak or K562 footprint. Yellow shading denotes the location of the ChIP-nexus peak (left) or K562 footprint (right). This illustrates the Fourier-based prior’s highlighting biologically relevant motifs outside of peak regions; the model trained without the prior, on the other hand, identifies these motifs more weakly/noisily, or not at all. Several mechanisms exist by which secondary motifs outside the central peak region can nonetheless assist TF binding within the peak region (e.g. through cofactors [13] or via 1D sliding [14]). A binary prediction model would be correct in identifying such motifs as being predictive of peak strength. Note that in contrast to binary models, profile models are less likely to detect such secondary motifs, as these motifs contribute to peak strength without contributing to the shape of the peak itself (the latter is primarily dictated by motifs lying within the central peak region).

**Figure S21:**
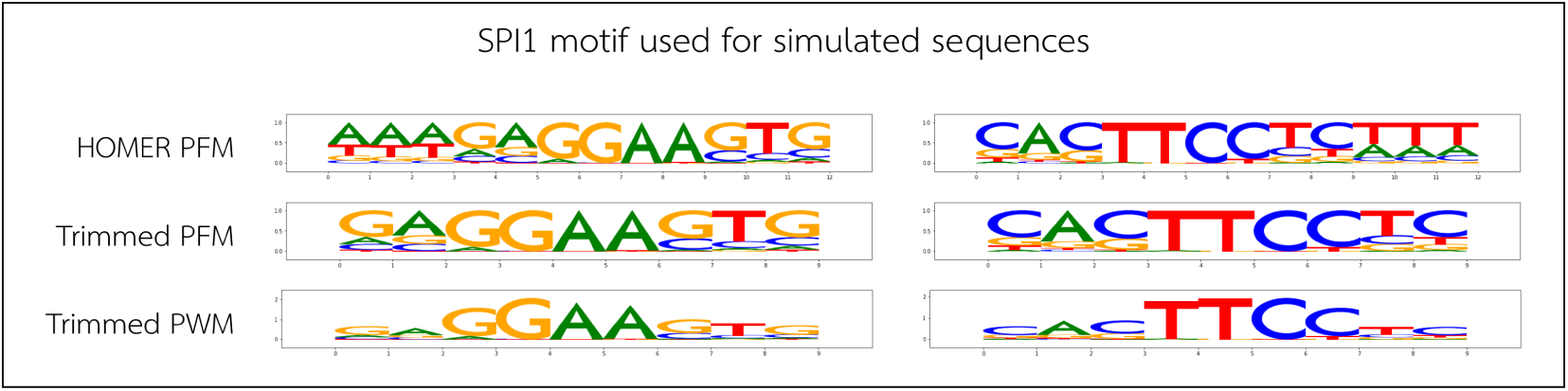
SPI1 motif used in constructing simulated sequences. We show the SPI1 motif (left) and its reverse complement (right) used to simulate SPI1-binding sequences for models trained on simulated data. Shown here are (from top to bottom): the Position Frequency Matrix (PFM) of the top motif identified by running HOMER 2 on IDR-thresholded peaks for SPI1 (Supplementary Methods Sec. 2.7); the trimmed motif PFM after removing flanks with low information content; and a PWM of the trimmed motif derived from weighting the PFM by information content.

**Figure S22:**
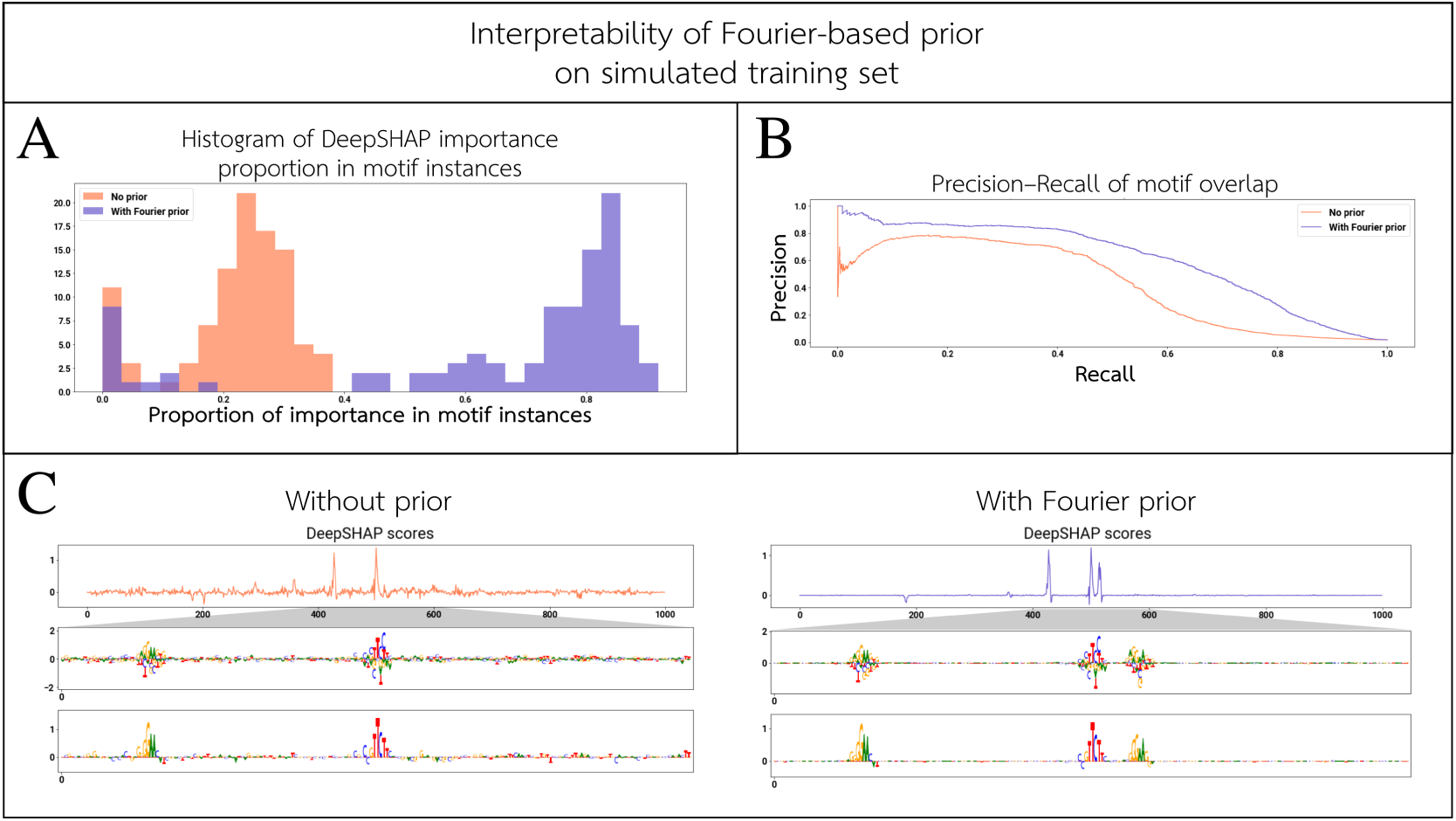
Attributions in simulated motif instances. We examine binary models trained to predict single-task SPI1 binding on simulated sequences. On a random sample of 100 motif-containing sequences, we compare models trained with versus without the Fourier-based prior by computing: **A)** the proportion of DeepSHAP importance overlying a motif instance; and **B)** the precision–recall of importance-ranked bases overlying motif instances. **C)** We also select an example sequence, and show the DeepSHAP attributions; the model trained with the Fourier-based prior cleanly highlights all three motif instances. Of the two zoomed-in attribution score tracks in Panel **C**, the top track represents the hypothetical importance scores (i.e. the importance that would be given to each of the four bases at each position, even if that base were not in the input sequence—see Supplementary Methods Sec. 3), and the bottom track represents the actual importance scores. See Supplementary Figure S21 for the SPI1 PWM used in the simulations. Note that the central motif instance is in the reverse-complement orientation.

**Figure S23:**
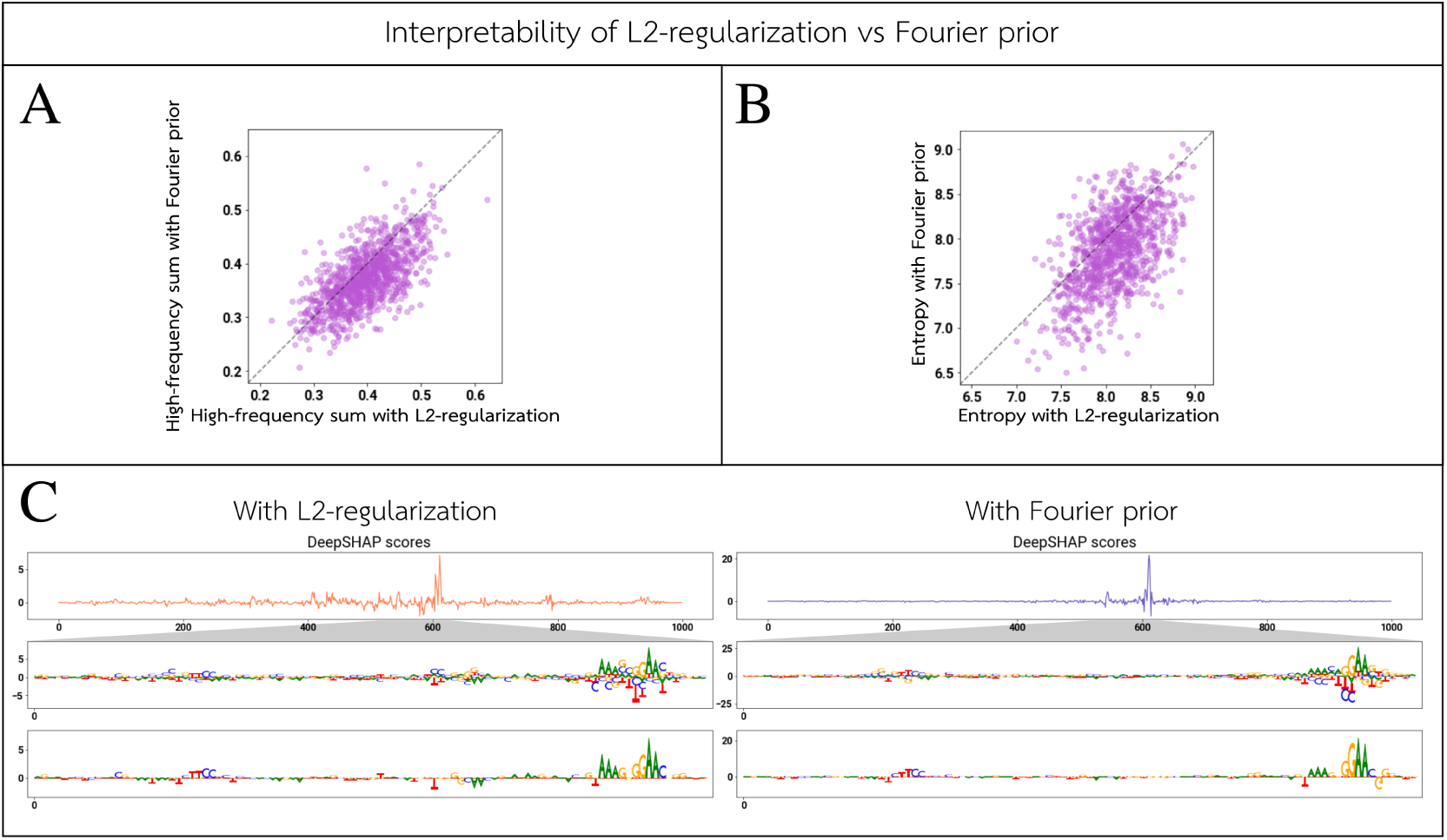
Signal-to-noise ratio of attributions with L2-regularization versus Fourier-based prior. For a SPI1 binary model, we show the DeepSHAP attributions for 1000 randomly selected peak sequences from the test set. We compare the sum of normalized high-frequency Fourier components and the Shannon entropy for each sequence, between a model trained with L2-regularization (i.e. weight decay) and a model trained with the Fourier-based prior. For a single selected test peak, we also display the value of the DeepSHAP importance for the bases present along the entire input sequence, as well as a zoomed-in view of a region close to the peaks summit. We also show the “hypothetical” attributions along each input sequence (i.e. the importance that would be given to each of the four bases at each position, even if that base were not in the input sequence—see Supplementary Methods Sec. 3).

**Figure S24:**
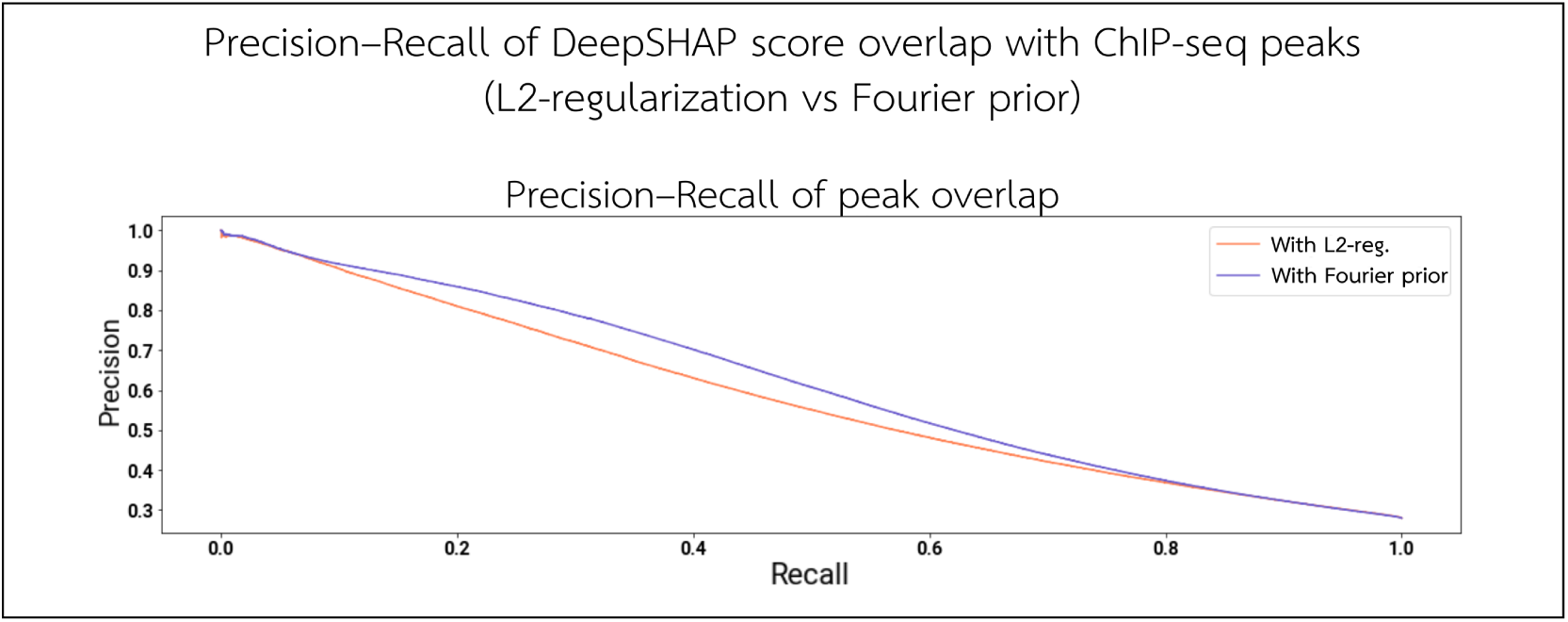
Precision–recall of important bases overlapping with ChIP-seq peaks (L2-regularization versus Fourier-based prior). For a SPI1 binary model, we sample 1000 peak sequences from the test set and compute the DeepSHAP attributions. We rank bases in descending order of total importance and generate a precision–recall curve by treating the set of bases that overlap a SPI1 ChIP-seq peak as “positives”. Shown here are the precision–recall curves for models trained with L2-regularization versus the Fourier-based prior.

**Figure S25:**
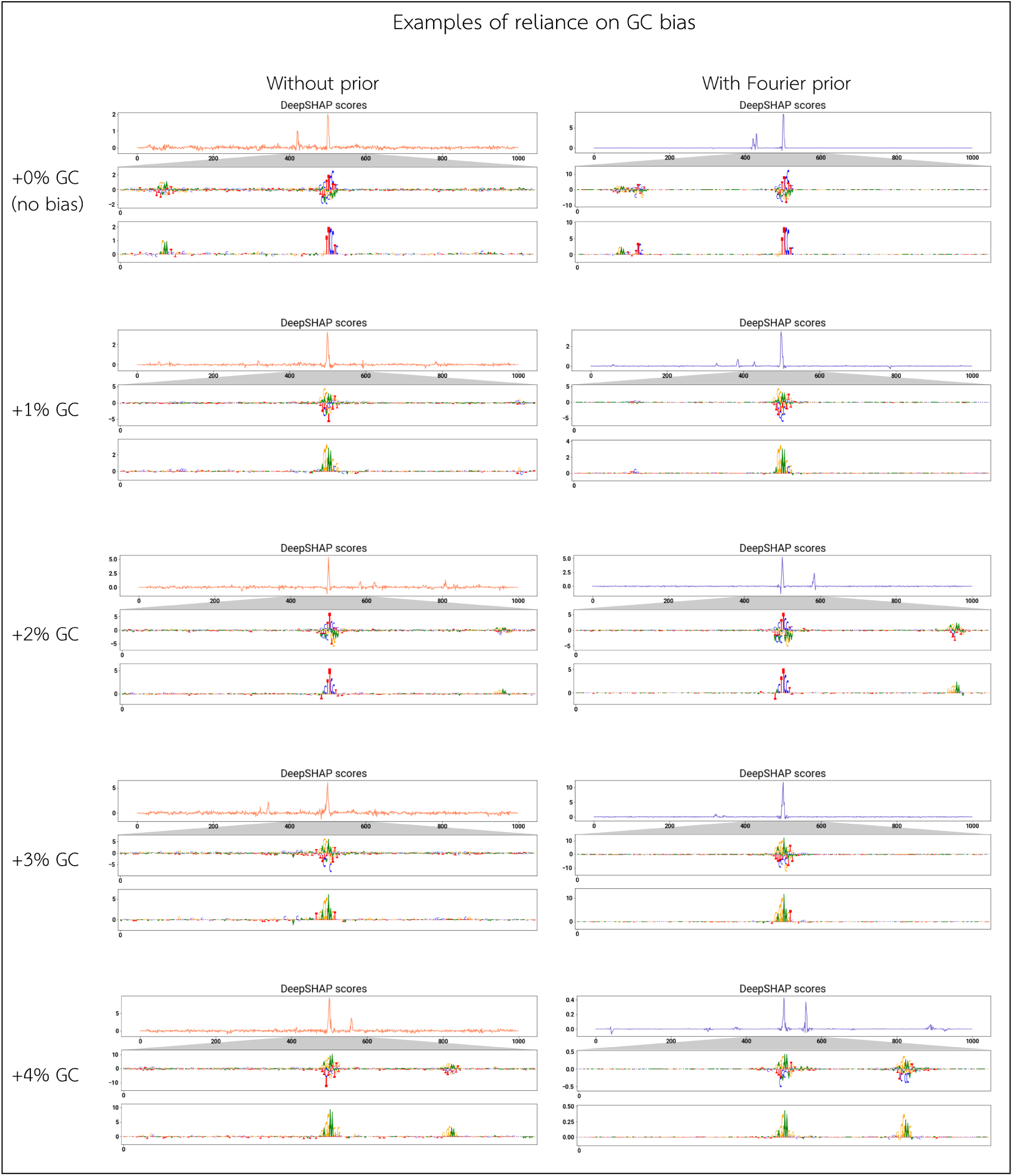
Specific examples of attributions under various levels of GC content. For each level of GC bias, we select a sampled input sequence and examine the DeepSHAP attributions. GC content ranges from +0% to +4%: a level of GC bias of +*x*% means the probability of G or C in the positive background sequences is (50 + *x*)%, while the negative sequence background has a G/C probability of 50% (i.e. no bias at all). We display the value of the DeepSHAP importance for the bases present along the entire input sequence, as well as a zoomed-in view of a region close to the peak summit. We also show the “hypothetical” attributions along each input sequence (i.e. the importance that would be given to each of the four bases at each position, even if that base were not in the input sequence—see Supplementary Methods Sec. 3).

## Supplementary Methods

### 1 Training data preparation

#### 1.1 SPI1 and GATA2 TF ChIP-seq

For these two transcription factors (TFs), we obtain data accessible through the ENCODE portal (https://encodeproject.org/) and processed using the ENCODE ChIP-seq pipeline [15, 16]. We select every TF ChIP-seq experiment with SPI1 or GATA2 as a target, satisfying the following conditions:

1. Experiment is of “released” status
2. Experiment has available unfiltered alignment BAMs aligned to hg38
3. Experiment has available BEDs of called peaks and IDR-filtered peaks aligned to hg38
4. Experiment has a matched control ChIP experiment with unfiltered alignment BAMs aligned to hg38
5. Cell type utilized in assay is not genetically modified

Based on these filtering criteria as of 4 Oct 2019, the ENCODE experiment IDs for SPI1 are ENCSR000BGQ, ENCSR000BUW, ENCSR000BIJ, ENCSR000BGW. The ENCODE experiment IDs for GATA2 are ENCSR000EVW, ENCSR000EWG, ENCSR000EYB. For each experiment, we obtain the unfiltered alignments, called peaks, and IDR-thresholded peaks, as available on the ENCODE portal (https://encodeproject.org/). For each TF, each experiment constitutes an output in the multi-task prediction models.

#### 1.2 K562 DNase-seq

We utilize a single DNase-seq experiment available through the ENCODE portal (https://encodeproject.org/) p: ENCSR000EOT [15, 16]. For this experiment, we obtain the unfiltered alignment BAMs, the set of called peaks that pass the ENCODE IDR filter, and the set of called peaks that did not pass the IDR filter. These files were computed using the ENCODE ATAC-seq pipeline [17].

Unlike TF ChIP-seq experiments, which have a matched control experiment, we perform bias correction by utilizing a control track that encodes the innate preferences of DNase [18]. We utilize the reads generated using this procedure, available as SRR1565781. These reads are processed into BigWigs using the ENCODE ATAC-seq pipeline into a single unstranded BigWig [17].

#### 1.3 Nanog/Oct4/Sox2 TF ChIP-seq

Nanog, Oct4, and Sox2 TF ChIP-seq experiments in mouse ESCs were performed by Avsec et al. [4]. We utilize the stranded read BigWig tracks and IDR-thresholded called peaks as prepared by the authors. This constitutes three different experiments (i.e. prediction tasks).

#### 1.4 Binary dataset preparation

The IDR-thresholded peaks are considered to be the high-confidence peaks. For the SPI1 and GATA2 TF ChIP-seq datasets, the 150,000 highest-scoring called peaks that do not overlap with an IDR-thresholded peak are considered ambiguous peaks, obtained using BEDtools v2.25.0 [19]. For the K562 DNase-seq dataset, the set of ambiguous peaks is generated by the ENCODE ATAC-seq pipeline [18]. For the Nanog/Oct4/Sox2 TF ChIP-seq datasets, there are no available ambiguous peak sets.

For all binary models, label generation consists of the entire hg38 genome split into 200 bp consecutive windows. A window is given a positive label if at least 100 bp of the window overlaps with a high-confidence peak, otherwise an ambiguous label if at least 100 bp of it overlaps with an ambiguous peak, and a negative label if neither of these two cases hold. For the TF ChIP-seq binary models, which have multiple tasks, the label generation procedure is performed for each task (i.e. each window will have an associated label for each task). These windows are then padded with 400 bp of context sequence on either side to form the final 1000 bp input to the network. This binary label generation was performed using seqdataloader [20].

#### 1.5 Profile dataset preparation

For the SPI1 and GATA2 TF ChIP-seq datasets and the K562 DNase-seq dataset, we first merge together the unfiltered reads of all biological replicates, and then filter them by keeping only reads with quality at least 30, and using samtools v1.2, with flag 780 [21]. This is nearly identical to the ENCODE TF ChIP-seq and ATAC-seq pipelines [15, 17], except this retains duplicate reads, which may be useful in the prediction of profiles.

Using bedGraphToBigWig v4 [22], we convert these reads into BigWigs of 5’ counts. For TF ChIP-seq datasets, the BigWigs are split into positive and negative strands, but for the K562 DNase dataset, the BigWig is unstranded. This gives each experiment a pair or a single BigWig track of 5’ counts.

The profile BigWigs for the Nanog/Oct4/Sox2 dataset were obtained from Avsec et al. [4] and are used as-is.

### 2 Model training

For both model architectures, we use a batch size of 128 and a learning rate of 0.001. We train for a maximum of 20 epochs with early stopping (requiring an improvement in the validation loss by at least 0.001 over the last 3 epoch deltas). Batch size, learning rate, and early stopping criteria were selected by choosing values in previous works on similar architectures, and verifying that validation performance was comparable.

We also utilize reverse complement augmentation in each batch, thereby effectively doubling the batch size.

Binary models learn a binary label from a 1 kb input sequence. Profile models learn a 1 kb profile from a 1346 bp input sequence.

Models were trained using PyTorch 1.3.0, on Google Cloud using n1-standard-8 instances, each with an NVIDIA Tesla P100.

#### 2.1 Training binary models

The binary model architecture consists of three consecutive 1D convolutions on the one-hot encoded DNA sequence. The convolutional layers have filter sizes of 15, 15, and 13, respectively (stride of 1). For TF ChIP-seq datasets, we use 64 filters in each layer; for K562 DNase-seq models, we use 256. The convolutional layers have ReLU activations and batch normalization. The result of the convolutions are fed into a max-pooling layer of size 40 and stride 40. The max-pooling output is fed into two consecutive fully connected layers of size 50 and 15, respectively, both with ReLU activations. Finally, the result is passed to a final dense layer that outputs a sigmoid prediction.

The loss function is binary cross-entropy, averaged across the different tasks. The cross-entropy loss for the positive and negative classes are averaged.

In each epoch, the network sees all genome bins where at least one task has a positive label. Each batch consists of an equal number genome bins where no task had a positive label, randomly subsampled at the beginning of each epoch.

#### 2.2 Training profile models

The profile model architecture, adapted from Avsec et al. [4], has a profile output and a total counts output for each task. The architecture consists of seven consecutive 1D dilated convolutions on the one-hot encoded DNA sequence. The first dilated convolutional layer has a filter size of 21, and the subsequent layers have filter sizes of 3 (all have stride of 1). For TF ChIP-seq datasets, we use 64 filters; for K562 DNase-seq models, we use 256. Dilation size is 1 for the first layer (i.e. no dilation), and increases by powers of two in subsequent layers. The dilated convolutional layers have ReLU activations. These dilated convolutional layers have summed residual connections, where the input to each layer is the sum of the outputs of all previous layers.

To compute the profile prediction, the last dilated convolution output is fed to another convolutional layer with kernel size 75 (stride 1) with no activation function. Finally, this result is then stacked with the control profile tracks, and fed to a final convolutional filter of size 1, such that the filter operates on one base of the logits and control profiles at a time. This constitutes the predicted profile logits.

To compute the total read count prediction, the last dilated convolution output is fed to a global average pooling layer, then to a dense layer. The result is concatenated with the total read counts in the control experiment, and fed through a final dense layer that predicts the log of the total read counts.

To compute the profile output and profile loss, the predicted profile logits are converted into probabilities by passing through a softmax along the profile prediction dimension. This gives the predicted profile shape. For each track, these probabilities, along with the true read counts over the region, define a multinomial distribution, where each base is a bucket. The profile loss is the log probability of seeing the distribution of true reads over this distribution, where the post-softmax predictions define the likelihood of a true read falling in that base. The results over all strands and tasks are averaged.

To compute the total counts loss, the predicted counts are treated as log counts, and the counts loss is simply a mean squared error of the log total counts, averaged over all strands and tasks (with a pseudocount of 1 for numerical stability).

When training, the profile loss and counts loss are given a weight of 1 and 20, respectively.

A positive example for a profile model is an input sequence and target profile track set centered at an IDR-thresholded peak summit (for any of the tasks). In each epoch, the network sees all IDR-thresholded peaks (aggregated over all tasks). Each batch consists of an equal number of non-peak sequences/profiles, sampled uniformly at random from the genome per batch. This implies that a sampled sequence intended for the negative set may overlap a peak, however unlikely it may be. The peak sequences are also randomly jittered up to 128 bp from the summit in either direction, to augment the set of positive examples. For the K562 profile models, however, we do not train with a negative set due to time efficiency, as we anecdotally found that the profiles with random jitters were sufficient to yield good performance.

Our profile models also utilize control profiles for bias correction. For the SPI1, GATA2, and K562 models, these are matched controls. For the Nanog/Oct4/Sox2 profile models, the control is identical and multiplexed across all tasks.

Our TF ChIP-seq profile models (i.e. for SPI1, GATA2, and Nanog/Oct4/Sox2) are stranded, and predicted and control profiles have positive and negative strands. Our K562 DNase-seq profile models, however, are unstranded, and the predicted and control profiles are summed across both strands.

#### 2.3 Fourier-based prior loss

In each batch during training, we only compute the Fourier-based prior loss for positive examples (i.e. binary examples where at least one task had a positive label, or profile examples originating from a peak). The Fourier-based prior loss is computed separately for each positive input sequence in the batch, and the loss is averaged over all positive input sequences.

The attributions *g*(*f, x*) are computed for each positive example, resulting in a vector of the same length *N* as the input sequence. In our models, *g*(*f, x*) is based on the element-wise product of the input one-hot-encoded sequence and the gradient of the output logits with respect to the input (as described in Shrikumar et al. [23]). This element-wise product is then summed over the base axis, yielding a single score for each position in a given sequence. For binary models, we simply multiply the input sequence and the gradient of the binary logits with respect to the input sequence. For profile models, analogous to Avsec et al. [4], we use the profile prediction logits (pre-softmax), weighted by the post-softmax probabilities, and summed across the entire profile; these logit gradients are then multiplied by the input sequence to yield *g*(*f, x*). For models with multiple tasks or strands, these logits are summed across these dimensions; thus, every positive training example will have a vector of attributions *g*(*f, x*) that is the same length as the input sequence.

To compute the attribution prior loss, We take the absolute value of these attributions, and smooth them with a Gaussian kernel *G* of width 1 standard deviation on both sides, with standard deviation equal to 3 base pairs. Let the resulting smoothed attribution vector be *g*^*s*^(*f, x*). Then we compute the discrete Fourier transform and recover the magnitudes *m* of the positive Fourier frequencies:

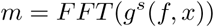

Note that *m* is a vector of length 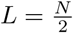. We discard the component corresponding to DC (i.e. the average value of the attributions) and *ℓ*_1_-normalize the magnitudes:

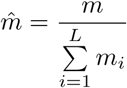

Finally, we compute the attribution prior loss as the sum of the high-frequency normalized magnitudes:

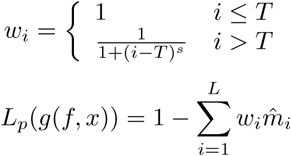

We utilize a soft cut-off, with *s* = 0.2. This cut-off was selected by visualizing the graph of 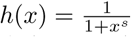 and selecting a reasonable *s* such that *h*(*x*) decays gracefully over a span of roughly 50 bp (Supplementary Figure S1).

For a signal of length *N*, a rectangular pulse of size *p* has a discrete Fourier transform of a sinc function with zeros every integer multiple of 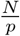. For binary models, the cut-off *T* is selected to be 150 (in terms of frequency index), and for profile models, the cut-off is 200. In both cases, this corresponds to a motif length (i.e. pulse) of 6–7 (the input sequences to the binary and profile models have different lengths, and hence a different frequency index corresponds to a pulse of the same size).

When training with the Fourier-based prior, the prior loss is weighted by 1 for binary models. For profile models, we train a model first without the Fourier-based prior, we take half of the validation loss (rounded to the nearest multiple of 5) after the model converges to be the weight of the prior loss (Supplementary Figure S4).

#### 2.4 Peak subsampling

In some of our downstream analyses, we train models with only a subset of the dataset (e.g. 1% of the training set). To subsample our dataset, we limit the set of IDR-thresholded peaks to only the top 1% by signal strength. We do this by ranking all IDR-thresholded peaks across all tasks in descending order by signal strength, remove duplicate peaks (i.e. perfect overlaps), and retain only the top 1% by count. This keeps only the 1% strongest/most confident peaks. For binary models, we recreate the training set from this new set of limited peaks. Negative bins are selected during training as usual. For profile models, we simply train with this smaller set of peaks as the positive set. Negative examples training are selected genome-wide, as usual.

For the Nanog/Oct4/Sox2 profile models trained on just 1% of the training data, we train for 80 epochs instead of the usual 20 (both with or without the Fourier-based prior).

#### 2.5 Training logistics

In all models, we reserve chr1 as the test set, reserve chr8 and chr10 as the validation set, and partition all other chromosomes in the training set (for hg38 or mm10). Evaluation of a model on the validation and test set proceeds identically to the training set in terms of the selection of positive and negative examples (i.e. we utilize all positive examples in validation/test, and an equally sized random sample of negative examples).

To evaluate binary model performance, we examine the validation loss, and the test set accuracy, auROC, and auPRC. These metrics are computed using balanced positive and negative classes. Because the auPRC is sensitive to class imbalance, we also compute an “estimated test auPRC”, which estimates the true auPRC on the full test set as if the negative examples were not subsampled to achieve a balanced set. We use this estimate rather than computing the auPRC on the full test set for computational efficiency. This estimate is calculated by artificially inflating the false positive rate, with the assumption that the negatives already present are representative of the full distribution of negatives.

To evaluate profile model performance, we examine the profile loss (i.e. the negative log-likelihood of the profiles).

For each dataset and model architecture, we train the model using 30 different random initializations. To compare a model trained with versus without the Fourier-based prior in our downstream analyses, we always compare the model with the best validation loss over all random initializations and epochs. For profile models, we compare the profile validation loss, rather than the aggregate loss of the profiles and counts outputs.

#### 2.6 L2-regularization models

We train SPI1 binary models with L2-regularization (without the Fourier-based prior), using an added L2-norm loss, consisting of the L2-norm of all trainable parameters in the network.

We tuned the L2-norm loss weight over 25 logarithmically sampled random weights from 10^−8^ to 10^2^, eventually settling on the optimal weight of 0.0001, which yielded the lowest validation loss. Using this optimal L2 loss weight, we trained 20 random initializations of SPI1 binary models. In all analyses performed, we used the model/epoch with the best validation loss.

#### 2.7 Models on simulated sequences

We train single-task SPI1 binary models with simulated DNA sequences for the positive and negative labels. We also only train for 5 epochs, where each epoch has the same number of positive and negative examples as a single-task SPI1 binary model trained on ENCSR000BGQ. All other training details are identical to the SPI1 binary models on real experimental data.

To create the input sequence for a negative label, we synthesize a sequence where every base is sampled independently and uniformly from A, C, G, and T. To create the input sequence for a positive label, we similarly sample a sequence from a uniform distribution of bases, and subsequently place a single instance of the SPI1 motif in the center. The placed motif is sampled from a PWM that was generated using HOMER 2 [24] on the set of IDR-thresholded peaks for SPI1, aggregated over all 4 ENCODE experiments (Supplementary Figure S21). Specifically, we used findMotifsGenome.pl in hg38 using the IDR-thresholded peaks, with a length of 12 and size of 200. We keep only the top motif. We then trim the motif by removing flanking regions with less than 20% of the information content of the base with the highest information content in the motif. Positive and negative sequences are sampled randomly at training-time every time a batch is generated. We use simdna to generate simulated positive and negative sequences [25].

We trained models with and without the Fourier-based prior over 3 random initializations each, and picked the model and epoch with the lowest validation loss for all downstream analyses.

### 3 DeepSHAP computation

To compute attributions (i.e. importance scores) for input sequences, we utilize DeepSHAP [11]. For binary models, we explain the binary prediction logits, summed across tasks. For profile models, we explain the profile prediction logits weighted by the final post-softmax probabilities, summed across the profile, and summed across tasks and strands.

Our baseline/reference set for DeepSHAP computation consists of 10 randomly shuffled versions of the input sequence, preserving dinucleotide frequencies. This choice of reference was recommended for genomic sequences in the DeepLIFT paper [7].

In subsequent sections, we use the term “hypothetical” DeepSHAP importance to refer to the estimated importance scores of an input sequence for all possible bases (i.e. the estimated DeepSHAP attributions for each base if it were present, hypothetically, at that location). The procedure for computing “hypothetical” importance scores can be found in Shrikumar et al. [6]. We use the term “actual” DeepSHAP importance to refer to the hypothetical importance multiplied by the input sequence (i.e. projected onto the bases that are actually in the sequence) and reduced to a single dimension along the input sequence by summing over the base identities at each position.

In order to make DeepSHAP work with PyTorch models and to easily produce “hypothetical” importance scores, we used a slightly modified version of the DeepSHAP library, available at https://github.com/atseng95/shap.

### 4 Signal-to-noise ratio of attributions

For each dataset, we select 1000 random positive examples from the test set and examine their interpretability. For profile models, we include a random jitter of ± 128 bp to avoid center-bias. We compute the “actual” DeepSHAP importance scores as described above for each sequence and quantify each sequence’s signal-to-noise ratio as the sum of the high-frequency normalized Fourier components, as well as the Shannon entropy. For simplicity, the sum of high-frequency normalized Fourier components follows the same definition as above, but with *s* = *∞* (i.e. no softness).

We use this same procedure to compare the attributions of our SPI1 binary models trained with L2-regularization versus the Fourier-based prior.

### 5 Motif discovery and motif calling

For our Nanog/Oct4/Sox2 models, we call motifs using TF-MoDISco v0.5.5.5 [6]. On the entire set of IDR-thresholded peaks in the test set, we compute DeepSHAP importance scores on the individual tasks for each TF, and run TF-MoDISco. We utilize a sliding window size of 21, flank size of 10, and seqlet FDR of 0.01. From the resulting motif clusters, we select only the motifs that have an average information content of at least 0.6 over some window of length 6. We then trim the motifs by removing flanking regions with less than 20% of the information content of the base with the highest information content in the motif. For binary models, due to their innate limitations in distinguishing primary motifs, we further filter motifs to have at least 750 supporting seqlets. For each motif, this gives us a PWM and a CWM (Contribution Weight Matrix). We also remove motifs corresponding to homopolymer repeats. The CWM consists of the aggregated “actual” DeepSHAP importance scores [4]. The background frequencies used to compute the PWM are 27% A or T, and 23% G or C.

To call motif instances, we select 1000 random positive examples from the test set. For profile models, we include a random jitter of ± 128 bp to avoid center-bias. For each motif, we compare it to every possible window in these sequences, and call a motif instance if:

1. The PWM match score is positive
2. The summed continuous Jaccard similarity (as defined below in Sec. 7) between the motif CWM and the underlying “actual” DeepSHAP scores of the sequence window is in the top 10% for that motif

We then rank the motif calls by the total DeepSHAP importance magnitude, summed across the instance, and compute precision–recall for which motif instances overlap with corresponding ChIP-nexus peaks by at least 1 base.

### 6 Performance on binary models

On our binary models, we compute accuracy, auROC, and auPRC on the test set, randomly subsampling negative bins to achieve balanced positives and negatives. We also compute an estimate of the auPRC on the imbalanced test set by artificially inflating the false positive rate to simulate auPRC computation on the entire test set. These are computed over all 30 random initializations for each dataset. The performance metrics of the best-performing model over the random initializations is recorded, with the best-performing model being the model with the lowest validation lost over all random initializations and epochs.

### 7 Stability of model learning

For each dataset, we select 100 random positive examples from the test set and examine the consistency of their “hypothetical” DeepSHAP importance across different models. For profile models, we include a random jitter of ±128 bp to avoid center-bias.

For each input sequence, we compute the similarity of “hypothetical” DeepSHAP importance from two models using the continuous Jaccard similarity metric [6]. The continuous Jaccard similarity is computed between the two sets of *ℓ*_1_-normalized “hypothetical” attribution scores for each base, and the resulting similarities are summed across the length of the sequence. For two *d*-vectors *u* and *v* (here, *d* = 4 for the 4 bases), the Jaccard similarity per base is computed as follows:

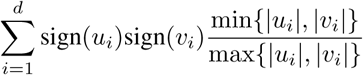

To quantify the similarity across all 30 random initializations of a model, for each of the sampled input sequences, we compute the pairwise continuous Jaccard similarity sum for each pair of models and average the scores across all 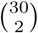 pairs. To quantify the similarity between models trained with all versus only 1% of the training data, we select the top 5 best-performing models when trained with all data, and the top 5 best-performing models when trained with only 1% of the training data, and we compute the pairwise continuous Jaccard similarity sum across all 5 × 5 pairs between the two conditions. The best-performing models are selected by picking the 5 models (5 different random initializations) in each condition with the best validation loss (or profile validation loss, for profile models).

### 8 Reliance on biologically relevant regions

For each dataset, we select 1000 random positive examples from the test set and examine the relationship between the “actual” DeepSHAP attributions and the underlying summits/peaks. For profile models, we include a random jitter of ±128 bp to avoid center-bias.

First, over each of the 1000 sampled sequences, we compute the Spearman correlation of the magnitude of the “actual” DeepSHAP importance at each base to the distance to the closest IDR-thresholded peak summit.

Next, over all 1000 sampled sequences in aggregate, we rank all bases in descending order by the magnitude of “actual” DeepSHAP importance, and ask whether the top bases overlie IDR-thresholded peaks. We use this to compute precision–recall curves, where thresholds are the “actual” DeepSHAP magnitude, and a positive is when a base overlies an IDR-thresholded peak.

We use this same procedure as above to compare the attributions of our SPI1 binary models trained with L2-regularization versus the Fourier-based prior.

For the K562 models and Nanog/Oct4/Sox2 models, we perform further analyses using orthogonal footprints or ChIP-nexus data, respectively. The K562 footprints are computed by Vierstra et al. [12]. From their published footprints, we aggregated the footprints of all K562 experiments using BEDtools merge [19]. For the Nanog/Oct4/Sox2 models, in addition to ChIP-seq experiments, Avsec et al. [4] also performed ChIP-nexus experiments, and we utilize their IDR-thresholded peaks from the ChIP-nexus experiments.

To compute the fraction of importance in a ChIP-nexus peak or footprint, we take the absolute value of the “actual” DeepSHAP importance of each of the sampled sequences, and calculate the proportion (out of the total sum across that sequence) within a ChIP-nexus peak or DNase footprint. We also repeat the rank-based analysis with footprints and ChIP-nexus peaks instead of ChIP-seq or DNase-seq peaks.

We also compute the motif clusters of the high-importance regions that the Fourier-based prior highlighted, but where the regions did not overlie a ChIP-nexus peak or a footprint. Over the same sample of 1000 test sequences used to compute the auPRC of ChIP-nexus peak or footprint overlap, we take the top 200000 most important bases (i.e. highest magnitude of DeepSHAP score) for which the base did not overlie a ChIP-nexus peak or a footprint. For each base, we expand to a centered 50 bp region, discarding overlaps (keeping the higher-ranked bases). For the resulting regions, we run the TF-MoDISco v0.5.5.5 [6] clustering algorithm only, clustering seqlets of length 30 into motifs.

To evaluate the ability of our models trained on simulated data to place importance only on motifs, we employ a similar procedure as above. We take a random sample of 100 simulated positive sequences (i.e. with placed motifs). For each sequence, we identify instances of the SPI1 motif by considering locations where the match score to the SPI1 PWM (i.e. the PWM used to construct the sequences) is over 0.9. The match score of a potential motif instance is computed as the sum of the entries in the PWM corresponding to the bases in the instance (i.e. total log-odds). Here, we used a uniform background (i.e. 25% A, C, T, or G) to compute the PWM. To compute the fraction of importance in the motifs, we use the same procedure as above to compute fraction of importance in a ChIP-nexus peak or footprint, but use motif instances instead of peaks/footprints. To compute the auPRC of base overlap with motif instances, we use the same procedure as above to compute precision–recall of base overlap with peaks/footprints, but use called motif instances instead.

### 9 GC content simulations

We train single-task binary SPI1 models on varying amounts of GC bias. The procedure for constructing simulated sequences is identical to the procedure described above for training single-task binary SPI1 models without GC bias. The only difference is that when we synthesize a positive sequence (i.e. with an inserted motif instance), the background sequence can have a higher amount of GC content. The negative sequence set always has sequences of equal GC and AT content (i.e. no bias). We train models on 5 different levels of GC content: +0%, +1%, +2%, +3%, and +4%. A level of GC bias of +*x*% means the probability of G or C in the positive background sequences is (50 + *x*)%, while the negative sequence background has a G/C probability of 50% (i.e. no bias at all).

We trained models with and without the Fourier-based prior over 3 random initializations each, and picked the model and epoch with the lowest validation loss for all downstream analyses.

To quantify how much a model is relying on GC content in the background, we sample 100 random positive sequences with the specified amount of GC bias, and compare the importance of A or T to the importance of G or C. For each sequence, we first mask out all instances of the SPI1 motif by ignoring any locations where the match score to the SPI1 PWM (i.e. total log-odds) is over 0.9. When scanning each simulated sequence for matches to the SPI1 motif, we computed the PWM using background frequencies that match the GC content of the sequence. We then consider the (signed) “hypothetical” DeepSHAP importance of A or T versus G or C, normalized by the maximum importance over the entire “hypothetical” importance track. We compute this product per base, and average over each input sequence.

## Notes

https://github.com/atseng95/att_priors

### Competing Interest Statement

The authors have declared no competing interest.

## References

[1] ENCODE Project Consortium et al. An integrated encyclopedia of dna elements in the human genome. Nature, 489(7414):57–74, 2012.

[2] Babak Alipanahi, Andrew Delong, Matthew T. Weirauch, and Brendan J. Frey. Predicting the sequence specificities of DNA- and RNA-binding proteins by deep learning. Nature Biotechnology, 33(8):831–838, aug 2015. ISSN 15461696. doi:10.1038/nbt.3300.

[3] David R. Kelley, Jasper Snoek, and John L. Rinn. Basset: Learning the regulatory code of the accessible genome with deep convolutional neural networks. Genome Research, 26(7):990–999, jul 2016. ISSN 15495469. doi:10.1101/gr.200535.115.

[4] Žiga Avsec, Melanie Weilert, Avanti Shrikumar, Amr Alexandari, Sabrina Krueger, Khyati Dalal, Robin Fropf, Charles McAnany, Julien Gagneur, Anshul Kundaje, and Julia Zeitlinger. Deep learning at base-resolution reveals motif syntax of the cis-regulatory code. bioRxiv, page 737981, aug 2019. doi:10.1101/737981. URL https://www.biorxiv.org/content/10.1101/737981v1.full.

[5] Jian Zhou and Olga G. Troyanskaya. Predicting effects of noncoding variants with deep learning-based sequence model. Nature Methods, 12(10):931–934, sep 2015. ISSN 15487105. doi:10.1038/nmeth.3547.

[6] Avanti Shrikumar, Katherine Tian, Anna Shcherbina, Žiga Avsec, Abhimanyu Banerjee, Mahfuza Sharmin, Surag Nair, and Anshul Kundaje. TF-MoDISco v0.4.2.2-alpha: Technical Note. oct 2018. URL http://arxiv.org/abs/1811.00416.

[7] Avanti Shrikumar, Peyton Greenside, and Anshul Kundaje. Learning Important Features Through Propagating Activation Differences, jul 2017. ISSN 1938-7228. URL http://proceedings.mlr.press/v70/shrikumar17a.html.

[8] Chiyuan Zhang, Samy Bengio, Moritz Hardt, Benjamin Recht, and Oriol Vinyals. Understanding deep learning requires rethinking generalization. 5th International Conference on Learning Representations, ICLR 2017 - Conference Track Proceedings, nov 2016. URL http://arxiv.org/abs/1611.03530.

[9] Gabriel Erion, Joseph D. Janizek, Pascal Sturmfels, Scott Lundberg, and Su-In Lee. Learning Explainable Models Using Attribution Priors. jun 2019. URL http://arxiv.org/abs/1906.10670.

[10] Andrew Slavin Ross, Michael C. Hughes, and Finale Doshi-Velez. Right for the Right Reasons: Training Differentiable Models by Constraining their Explanations. mar 2017. URL http://arxiv.org/abs/1703.03717.

[11] Scott M Lundberg and Su-In Lee. A Unified Approach to Interpreting Model Predictions. In I Guyon, U V Luxburg, S Bengio, H Wallach, R Fergus, S Vishwanathan, and R Garnett, editors, Advances in Neural Information Processing Systems 30, pages 4765–4774. Curran Associates, Inc., 2017. URL http://papers.nips.cc/paper/7062-a-unified-approach-to-interpreting-model-predictions.pdf.

[12] Jeff Vierstra, John Lazar, Richard Sandstrom, Jessica Halow, Kristen Lee, Daniel Bates, Morgan Diegel, Douglas Dunn, Fidencio Neri, Eric Haugen, Eric Rynes, Alex Reynolds, Jemma Nelson, Audra Johnson, Mark Frerker, Michael Buckley, Rajinder Kaul, Wouter Meuleman, and John A. Stamatoyannopoulos. Global reference mapping and dynamics of human transcription factor footprints. bioRxiv, page 2020.01.31.927798, feb 2020. doi:10.1101/2020.01.31.927798.

[13] Samuel A Lambert, Arttu Jolma, Laura F Campitelli, Pratyush K Das, Yimeng Yin, Mihai Albu, Xiaoting Chen, Jussi Taipale, Timothy R Hughes, and Matthew T Weirauch. The human transcription factors. Cell, 172(4):650–665, February 2018.

[14] David M Suter. Transcription factors and DNA play hide and seek. Trends Cell Biol., 30(6):491–500, June 2020.

[15] Ian Dunham, Anshul Kundaje, Shelley F. Aldred, …, and Lucas Lochovsky. An integrated encyclopedia of DNA elements in the human genome. Nature, 489(7414):57–74, sep 2012. ISSN 14764687. doi:10.1038/nature11247.

[16] Carrie A Davis, Benjamin C Hitz, Cricket A Sloan, Esther T Chan, Jean M Davidson, Idan Gabdank, Jason A Hilton, Kriti Jain, Ulugbek K Baymuradov, Aditi K Narayanan, Kathrina C Onate, Keenan Graham, Stuart R Miyasato, Timothy R Dreszer, J Seth Strattan, Otto Jolanki, Forrest Y Tanaka, and J Michael Cherry. The Encyclopedia of DNA elements (ENCODE): data portal update. Nucleic Acids Research, 46(D1):D794–D801, 2017. ISSN 0305-1048. doi:10.1093/nar/gkx1081. URL https://doi.org/10.1093/nar/gkx1081.

[17] Jinwook Lee, Daniel Kim, Grey Cristoforo, Chuan-Sheng Foo, Chris Probert, Nathan Beley, and Anshul Kundaje. ENCODE ATAC-seq pipeline. dec 2019. doi:10.5281/ZENODO.3564806.

[18] Galip Gürkan Yardimci, Christopher L. Frank, Gregory E. Crawford, and Uwe Ohler. Explicit DNase sequence bias modeling enables high-resolution transcription factor footprint detection. Nucleic Acids Research, 2014. ISSN 13624962. doi:10.1093/nar/gku810.

[19] Aaron R. Quinlan and Ira M. Hall. BEDTools: A flexible suite of utilities for comparing genomic features. Bioinformatics, 2010. ISSN 13674803. doi:10.1093/bioinformatics/btq033.

[20] Annashcherbina, Av Shrikumar, and Soumya Kundu. kundajelab/seqdataloader: v0.2. apr 2020. doi:10.5281/ZENODO.3771365.

[21] Heng Li, Bob Handsaker, Alec Wysoker, Tim Fennell, Jue Ruan, Nils Homer, Gabor Marth, Goncalo Abecasis, and Richard Durbin. The Sequence Alignment/Map format and SAMtools. Bioinformatics, 2009. ISSN 13674803. doi:10.1093/bioinformatics/btp352.

[22] W. J. Kent, A. S. Zweig, G. Barber, A. S. Hinrichs, and D. Karolchik. BigWig and BigBed: Enabling browsing of large distributed datasets. Bioinformatics, 2010. ISSN 13674803. doi:10.1093/bioinformatics/btq351.

[23] Avanti Shrikumar, Peyton Greenside, Anna Shcherbina, and Anshul Kundaje. Not just a black box: Learning important features through propagating activation differences. May 2016.

[24] Sven Heinz, Christopher Benner, Nathanael Spann, Eric Bertolino, Yin C. Lin, Peter Laslo, Jason X. Cheng, Cornelis Murre, Harinder Singh, and Christopher K. Glass. Simple Combinations of Lineage-Determining Transcription Factors Prime cis-Regulatory Elements Required for Macrophage and B Cell Identities. Molecular Cell, 2010. ISSN 10972765. doi:10.1016/j.molcel.2010.05.004.

[25] Av Shrikumar, Johnny Israeli, Karl Leswing, Pgreenside, Bharath Ramsundar, Wainberg, and BenjaminrBond. kundajelab/simdna: Bugfix for simdata loading from non-gzipped files. jun 2019. doi:10.5281/ZENODO.3258813.

